# Genetic investigation of GPI anchored Bd37 orthologs in *Babesia divergens* group and use of recombinant protein for ecological survey in deer

**DOI:** 10.1101/2024.03.19.585777

**Authors:** Aya Zamoto-Niikura, Katsuro Hagiwara, Koichi Imaoka, Shigeru Morikawa

## Abstract

The Glycosylphosphatidylinositol (GPI) anchored protein group has great potential as an excellent immunodiagnostic marker, because of its high expression and necessity for parasite survival. *Babesia divergens* /*B. capreoli* group includes parasites with confirmed or possible zoonotic potential to cause human babesiosis. In this study, we investigated ortholog of Bd37, a GPI-anchored major merozoite surface protein of *B. divergens* sensu stricto, in the Asia lineage of the *B. divergens* /*B. capreoli* group. From two genomic isolates from sporozoites/sporoblasts, three *Bd37* gene variants, namely *Bd37 JP-A*, *JP-B,* and *JP-C,* were isolated with 62.3% −64.1% amino acid sequences identity. Discriminative blood direct PCR revealed that *JP-A* was exclusively encoded in all parasites infecting wild sika deer examined (n=22). While *JP-B* and *JP-C* genes were randomly detected in 12 and 11 specimens, respectively. Recombinant JP-A-based ELISA showed an overall positive rate of 13.9% in deer in Japan from north (Hokkaido) to south (Kyushu islands) (24 prefectures, n=360). This positive rate was twice as high as that examined by *18S rRNA*-based PCR (6.8%). Antibodies against recombinant JP-B and JP-C were also evident in the deer. This study demonstrated that the presence of three orthologs in the Bd37 gene family in Asia lineage and identified JP- A as an informative marker for serological surveys in Japan. This is the first report that diagnostic antigen of *Babesia* parasite was identified by a comprehensive analysis of genetic polymorphisms from a various developmental stage in host and vector…

**Importance:** *Babesia divergens* Asia lineage in *B. divergens*/ *B. capreoli* group is a parasite closely related to zoonotic pathogen *B. divergens* sensu strict (EU lineage) and *Babesia* sp. MO1(US-lineage). Large scale serodiagnostic system for this group has not been established. As the nature of the parasite’s antigenic differentiation to escaping from immunological attack in the host, investigation of diagnostic markers should consider such antigenic diversity inherited (circulating) in the population. We focused on the Glycosylphosphatidylinositol (GPI) anchor protein, Bd37, a major surface protein of the EU lineage, and investigated Asia lineage infecting sika deer and taiga tick in Japan. Three Bd37 ortholog genes (JP-A, JP-B, and JP-C) were isolated from the tick and deer, though only *JP-A* gene was exclusively encoded in the parasite’s genomes (n=36). In spite of sequence polymorphism in the N-terminal region, the antibody raised against the representative recombinant antigen, rJP-A2, reacted to various JP- A proteins. rJP-A2-based ELISA system revealed a positive rate in wild sika deer was 13.9% which is two times higher than that examined by genetic examination (PCR). GPI-anchored proteins are densely expressed and required for parasite survival. We showed GPI proteins including Bd37 and its ortholog are potentially excellent immunodiagnostic markers for emerging and growing human babesiosis.

## Introduction

Research and management of wildlife diseases is becoming increasingly important. Underlying this trend is the recognition that emerging zoonotic diseases are on the rise and that wildlife can be an important factor in controlling infectious diseases in humans, livestock, and companion animals. In particular, the unique characteristics of wild cervid, as a population, present public health challenges. Gibier is concerned about the direct effects on humans, such as the Hepatitis E virus and *Toxoplasma* infections. (Takahashi et al. 2022, Dubey et al. 2020). Additionally, cervids are ecologically and phylogenetically closely related to livestock, including bovine and swine and they are at risk of transmitting pathogens (such as brucellosis, bovine tuberculosis, wasting disease, and epizootic hemorrhagic disease, EHD)(Conner et al. 2008, Abrantes and Vieira-Pinto 2023, Forzan et al. 2017). Cervids also have an indirect effect due to major blood-sucking sources of ticks. Ticks transmit many tick-borne diseases to humans including Japanese spotted fever and Lyme disease. Thus, expansion of deer habitat over human living areas threatens controlling tick-borne infectious diseases (Tabara et al. 2019, Eisen et al. 2023).

Many of the genotypes and serotypes found in cervid have zoonotic potential(Matei et al. 2019, Dubey et al. 2020). Especially in *Piroplasmida*, data are accumulating that the host specificity is not as strict as previously thought. For example, the deer protozoan *B. capreoli* and the dog protozoan *B. canis* have been detected in Italian horses (Zanet et al. 2017), and conversely, the horse protozoan *Theileria equi* infects dogs (Hegab et al. 2023, Fritz 2010, Beck et al. 2009, Criado-Fornelio et al. 2003). It is noted that the selection of marker genes and the sequencing area of analysis vary in each study, so care must be taken when interpreting them. Hrazdilova et al. analyzed almost full-length *18S rRNA* and *cytochrome oxidase subunit 1 (cox1)* gene sequences and detected as many as four *Babesia* spp., *B. divergens*, *B. capreoli*, “deer clade” (*Babesia* sp. CH1) and *Babesia* sp. EU1 (*B. venatorum*) in deer in Chez (Hrazdilová et al. 2020).

Previously we found novel *Babesia divergens*-like sequences from deer in Japan and showed monophyletic clade formed by *B. divergens* in Europe (*B. divergens* sensu stricto), *B. divergens*-like in the United States and Japan and *B. capleoli* (*B. divergens/B. capreoli* group) by *18S RNA* and *beta-tubulin* gene-based phylogenetic trees. In the study, the 4 lineages were named EU lineage, US lineage, Asia lineage, and *B. capleoli* lineage, respectively (Zamoto-Niikura et al. 2018). EU lineage or *B. divergens* sensu stricto, is the main causal agent of human babesiosis in European countries. Since the first case was reported in 1957 in a splenectomized Yugoslavian farmer, approximately 40 human cases have been reported from Europe and attributed to *B. divergens* infection, although not all cases were diagnosed molecularly. *Ixodes ricinus*, the most important tick species in public health, and bovine are the principal vector and reservoir for this parasite, respectively. A nationwide serological survey using whole parasite as antigen in indirect fluorescence assay (IFA) revealed positive existence of antibodies against *B. divergens* was evident in blood donors in Austria (2.1%) (Sonnleitner et al. 2014), in tick-exposed patients in Germany (4.9%) (Hunfeld et al. 2002) and Belgium (33.2%) (Lempereur 2015), and in forestry workers in France (0.1%) (Rigaud et al. 2016), suggesting widely occurred subclinical infections in healthy persons.

The US lineage, or *B. divergens-*like MO1, emerged in 2002 in a patient in Kentucky, who had undergone splenectomy. Since then, 5 severe cases including a possible transfusion transmission have been reported in the United States (Burgess et al. 2017). Very recently, a *Babesia* sp. closely related to the MO1 (*Babesia* sp. FR1) emerged in a splenectomized patient in France (Bonsergent 2021), suggesting this lineage is distributed widely from North America to the Eurasian continent. *B. capleoli* (*B. capreoli* lineage of *B. divergens* group, in this study) have long been recognized as a parasite of cervids. While the retrospective study in northwestern China, Gansu province, revealed parasites in undiagnosed patients (Wang et al. 2019) had identical *18S rRNA* sequence (MK256977) to those of the *B. capleoli* lineage from reindeer (*Rangifer tarandus*) in Germany (KM657248) and roe deer (*Capreolus capreolus*) in France (FJ 944827). *B. divergens* Asia-lineage is maintained between wild sika deer (*Cervus nippon*) and *I. persulcatus* in the eastern part of Japan (Zamoto-Niikura et al. 2014, 2018, 2020)*. I. persulcatus* is the most common species causing tick bites and transmits many tick-borne diseases including Tick-borne encephalitis (Yoshii et al. 2017), Lyme diseases and relapsing fever (Fukunaga et al. 1995).

Glycosylphosphatidylinositol-anchored proteins (GPI-AP) are a group of membrane proteins anchored to the outer cell membrane via GPI, and widely expressed in eukaryotic cells including protozoan parasites (Guha-Niyogi et al. 2001) (Komath et al. 2022). The protozoan GPIs are distinguishable from mammalian GPIs by distinct side chains and are expressed about one hundred times more GPI-AP per cell than those of mammalian cells (Debierre-Grockiego 2010). GPI-APs have various biological functions. For example, *Trypanosoma brucei* expressed two GPI-APs on the cell surface in their blood stage, as transferrin receptor (TfR) and variant surface glycoprotein (VSG). TfR is a heterodimeric protein composed of GPI-anchored ESAG6 and ESAG7 and localizes in the flagellar pocket. (Kabiri and Steverding 2021). VSG is a homodimeric protein that coats the cell surface and allows the parasite to evade the humoral immune system and the host’s complement attack. Antigenically distinct VSGs are expressed by switching monoallelic selection from a repertoire of over 1500 genes. (Gadelha et al. 2011, Hertz-Fowler et al. 2008).

The EU lineage and *B. capleoli* (*B. capleoli* lineage) in the *B. divergens/ B. capleoli* express GPI-APs, Bd37 and its ortholog Bcp37/41, respectively. The Bd37 was initially identified as a soluble immunodominant antigen of 37kDa, which conferred protective immunity against homologous lethal challenges with Rouen87 human isolate (Gorenflot et al. 1990). The biochemical characters of the Bd37 include the presence of an N-terminal signal peptide, a C-terminal GPI-anchoring sequence, a disulfide bond bridging the N and C-terminal parts of the protein via cysteines, and charged residues involved in salt bridges (Delbecq et al. 2002). The *Bd37* sequences vary among different isolates, in particular, the N-terminal region (after signal peptide cleavage site to valine 80) is most polymorphic.

Nevertheless, a recombinant Bd37 of the Rouen87 conferred complete protection against lethal heterologous challenges with 5 different strains, which represented five major polymorphic groups identified by PCR-RFLP (Hadj-Kaddour 2007). Sequence polymorphism among isolates was also observed in *Bcp37/41* of the *B. capreoli*. Sequence identity between 40.8 k Da (clone 2801 F10) and 44 amino acid residues deleted 36.8 kDa (clone 2770 F6) is 50.3% (Sun et al. 2011). In all cases, the Bd37 and Bcp37/41 are considered to be expressed as single molecules from respective isolates.

The serological examination is widely used for diagnosis, screening and epidemiological surveys. Indirect fluorescence assay (IFA) by using whole parasites as antigens is a widely used method to detect antibodies against Babesial infection. The IFA antigen slide is prepared by spreading erythrocytes infected with *Babesia* isolate. This method can be useful for diagnosing the infection when a patient develops specific clinical signs for human babesiosis (Gonzalez et al. 2015). Since cross-reactivity among closely related species in IFA has been clearly shown with *B. divergens* (Rouen 1987 strain), *B. capreoli* and emerging zoonotic piroplasm, *B. venatorum* (Bastian et al. 2012), another serological tool is required to specifically detect *B. divergens* group infection and to conduct large scale epidemiological survey (Gabrielli et al. 2012).

Due to the lack of biological isolate of *B. divergens* Asia lineage, we thought the Bd37 ortholog in the lineage might be the candidate for diagnostic antigen. We attempted to isolate the genes from parasites naturally infected in ticks and deer’s erythrocytes. This direct investigation could lead us to discover the unique composition of the *Bd37* family in the individual parasites, and a establish serodiagnostic marker for detecting *B. divergens* Asia-lineage infection.

## Materials and methods

### DNA, RNA and cDNA preparation

The genomic isolates of *B. divergens* Asia-lineage, IpSG13-13-1 and IpSG10 (table 1) from *Ixodes persulcatus* ticks, were prepared in our previous study (Zamoto-Niikura 2018) and used for primely isolation of *Bd37* gene sequence. Briefly, the genomic DNA was isolated from activated salivary glands of ticks, which was fed on laboratory animals for 3 days. For RNA extraction, the salivary glands kept in −80 ℃ was used. Note that IpSG10 RNA was not available for this experiment since the salivary gland was not remained in freezer. Total RNA was extracted by using Isogen (Nippon Gene Co., Ltd.). Contamination of genomic DNA in the total RNA was assessed by nested PCR. Complement DNA was synthesized from the total RNA using *Bd37* specific primer, Bd37-TAA, which contains stop codon and upstream sequence (supplement table1) (SSC III reverse transcriptase, Invitrogen). Genomic DNA of MRNK strain (Zamoto-Niikura 2014) (table 1) was used for isolation of *Bd37* sequence as reference of *B. divergens* EU lineage.

**Table 1.** Materials.

| Material | *B. divergens* genetic isolate | Origin | Country | Reference |
| --- | --- | --- | --- | --- |
| gDNA, mRNA | IpSG13-13-1 | *Ixodes persulcatus* (field collected), salivary glands | Japan | Zamoto-Niikura 2018 |
| gDNA | IpSG10 | *Ixodes persulcatus* (field collected), salivary glands | Japan | Zamoto-Niikura 2018 |
| gDNA | MRNK | *Bos taurus* (maintained in gerbil) | Ireland | Zamoto-Niikura 2014 |

### PCR cloning and sequencing

Full length of *Bd37* sequence (app. 1kb), except for stop codon, was amplified by ExTaq polymerase (Takara Bio) using the primers, Bd37-ATG and Bd37-TAA removed (supplement table1), according to the manufacture’s protocol. The resulted PCR amplicons were purified (QIAquick PCR Purification Kit, Qiagen) and ligated into pBAD/Thio-TOPO plasmid vector (Thermo Fisher Scientific). The plasmids were transformed into TOP10 competent cells (Thermo Fisher Scientific). Transformants were screened by colony PCR using Bd37 internal primer Bd37-SQF1 (supplement table1) and vector specific primer pBAD-Reverse (Thermo Fisher scientific). Plasmids were purified by using Spin Miniprep Kit (Qiagen). Sequencing (Eurofins Genomics) was performed by using vector primer, Trx-Forward and pBAD-Reverse (Thermo Fisher Scientific). As a control, pBAD/Thio-TOPO plasmid vector carrying full length of Venus (YFP variant) fluorescent protein gene was generated accordingly (Zamoto-Niikura 2009).

### Analysis of *Bd37* gene sequences

DNA sequences were aligned with each other and with closely related sequences available in GenBank by ClustalW implemented in BioEdit7.2.5 (Hall 1999). DnaSP version 6 (Rozas et al. 2017) was used to determine the number of segregating sites (S), number of haplotype (H), haplotype diversity (Hd) and observed nucleotide diversity per site (π). The dN/dS (ratio of non-synonymous substitution / synonymous substitution) value was used as measure of evolutionary pressures on protein-coding regions by using Nei-Gojobori method with Jukes and Cantor calculate. A ratio dN/dS > 1 results when changes in the protein sequence are favored by natural selection (evidence of positive selection), while a ratio < 1 is expected if natural selection suppresses protein changes (evidence of negative selection). A dN/dS ratio equal to 1 represents a situation of neutral evolution. A two tailed Z-test was performed on the difference between Ds and Dn by MEGA-X 10.0.5 (Kumar et al. 2018).

### Analysis of Bd37 protein sequence

The putative protein sequences were aligned with each other and with closely related sequences available in GenBank by clustalW implemented in BioEdit7.2.5. Phylogenetic analysis was performed by the neighbor-joining (NJ) method with 1,000 bootstrap replicates in MEGA-X 10.0.5. Putative signal peptide and GPI-anchor were searched by SignalP-4.1 (http://www.cbs.dtu.dk/services/SignalP/), Phobius (http://phobius.sbc.su.se/) and PredGPI, (http://gpcr.biocomp.unibo.it/predgpi/pred.htm).

### Discriminative PCR

To detect and discriminate the *Bd37* gene sequence by the sequence types (JP-A, JP-B, JP-C and EU-A1 and EU-A2) (table 2), PCR primers specific for each type were designed. Primers and their specificity were listed in supplement table 1. It is noted that the specific primer JP-B/EU-A2 F2 anneals to both JP-B and EU-A2, since appropriate site could not be identified. The specificity of the designed primers was examined by using plasmid clones carrying either of JP-A, JP-B, JP-C, EU-A1 or EU-A2 type sequence (fig 3). In this test, the plasmids of 1×10^8^ copies in a PCR mixture were used. For examining sensitivity of the nested PCR, 1^st^ PCR was performed using universal primers Bd37-ATG and Bd37-TAA (supplement table 1) and 1×10^1^, 10^2^ or 10^3^ copies of plasmids in a PCR mixture (20μl). One micro little of the product was used for 2nd PCR in which either of the specific primers (forward) and universal primer Bd37-R1 (reverse) (supplement table 1) were used. Ex taq polymerase (Takara bio) was used according the manufacture’s instruction.

**Table 2.** Number of plasmid clones carrying various Bd37 sequence types.

| Isolate | Material | No. sequenced | Bd37 sequence type |  |  |  |  |  |  |
| --- | --- | --- | --- | --- | --- | --- | --- | --- | --- |
|  |  |  | JP-A |  |  | JP-B | JP-C | EU-A |  |
|  |  |  | 1 | 2 | 3 |  |  | 1 | 2 |
| IpSG13-13-1 | genome | 10 | 5 | 2 | 0 | 3 | 0 | 0 | 0 |
|  | cDNA | 10 | 3 | 0 | 0 | 7 | 0 | 0 | 0 |
| IpSG10 | genome | 10 | 0 | 3 | 4 | 0 | 3 | 0 | 0 |
| MRNK | genome | 10 | 0 | 0 | 0 | 0 | 0 | 6 | 4 |
| Total |  | 40 | 8 | 5 | 4 | 10 | 3 | 6 | 4 |

### Deer and tick samples used for the discriminative PCR

Erythrocytes of wild sika deer (*Cervus nippon*) kept in –80°C were used for blood direct PCR. Hunting, sample collection and screening of *B. divergens* infection were performed in our previous study. (Zamoto-Niikura 2014, 2020). *B. divergens* genomic DNAs extracted from host seeking *I. persulcatus* ticks (whole body) or additional salivary glands, IpSG 17-6-3, IpSG 14-12-2 were prepared in our previous study (Zamoto-Niikura 2018).

### Blood direct PCR and sequencing

To screen the presence of *Bd37* sequence in *B. divergens*-positive wild deer, frozen erythrocytes were melted and directly used as template (Phusion Blood direct PCR kit, Fisher Scientific) as described previously (Zamoto-Niikura 2020). Direct sequence was performed by using inner primers Bd37-SQF1 and Bd37-SQR1 (supplement table 1). Quality of base-calling was checked by Quality score in Sequencher (GeneCode), then chromatograph was manually checked in case of low-quality score (below 40%). When PCR amplicons were directly sequenced and overlapped peaks in chromatograph were observed, the corresponding residues were collected manually according to IUPAC nucleotide code. The threshold of detecting the minor population was 20%, which was calculated by comparing chromatograph of direct sequencing of PCR amplicon and sequencing of plasmids of PCR clones.

### Expression of recombinant Bd37 proteins

Recombinant Bd37 (rBd37) protein was expressed as thioredoxin fusion protein from the pBAD/Thio-TOPO vector (Thermo Fisher Scientific), which encodes thioredoxin and histidine-tag at N-terminus and C-terminus of multi cloning site, respectively. These additional sequences increase the size of expressed recombinant protein by 13 kDa and 3 kDa, respectively. Protein expression was induced with 0.2% - 2.0% of arabinose at 25 °C for 6 hours. Recombinant Bd37 and Venus (control) proteins were purified by using ProBond Purification system (Thermo Fisher Scientific) according to the manufacture’s instruction. Briefly, cells were harvested by centrifugation and lysed by sonication in native buffer including Complete Mini (Roche) and Lysozyme (Sigma Aldrich) (50mg/ml) on ice. Recombinant proteins were purified by nickel-chelating resin and eluted under native condition. The eluted fractions were concentrated by filter (Amicon Ultracel-30K, Merk Millipore) and dialyzed against TNE buffer (20mM Tris-HCl pH8.0, 0.5M NaCl, 0.1mM EDTA) at 4 °C. Protein concentration was measured using Pierce BCA Protein Assay Kit (Thermo Fisher Scientific). The purified rBd37 in TNE including sodium azide (0.02%) and DTT (0.2mM) were stored at 4 °C.

### Anti-sera against recombinant Bd37 protein

The purified rBd37 were dialyzed against PBS (-). Rabbit was injected with 200 μg of the rBd37 or rVenus in an equal volume of TiterMax Gold (TiterMax USA, Inc.) subcutaneously. Three weeks following primary immunization, same amount of antigen emulsified in TiterMax Gold was injected as booster. Whole blood was harvested after 7 weeks post initial immunization and serum was separated by centrifugation.

### SDS-PAGE and western blot

One hundred micro gram of the purified rBd37 or rVenus was loaded onto acrylamide gel (e-PAGEL 10-20% gradient, ATTO) with protein marker (Blue star, Nippon gene). After electrophoresis, separated proteins on the gel were stained with Coomassie Blue/ethanol/acetic acid and destained by methanol/acetic acid. For western blot, the separated proteins were transferred onto Immobilon-P PVDF membrane (Merck Millipore). The membrane was blocked with 10% skim milk in Tris Buffered Saline, with Tween 20, pH 8.0, and reacted with mouse anti-Thioredxin antibodies (MBL). Alkaline phosphatase conjugated goat anti-mouse IgG antibody (Sigma) was used as second antibody and proteins on the membrane were visualized using BCIP/NBT substrate (Sigma). When deer plasma specimens were used as primary antibody, the plasma was diluted by 1:50 with ImmunoBlock (DS Pharma Biomedical Co. Japan). ProteinA/G-Calf Intestinal Alkaline phosphatase conjugated (Thermo Fisher Scientific) was used as 2ndary antibody. Plasma specimens of wild sika deer were randomly chosen by prefecture from the sample collected in previous study (Zamoto-Niikura 2014, 2020).

### ELISA

Recombinant proteins were diluted with Bicarbonate/carbonate coating buffer (100mM) at 0.2 ug/mL and 100μL/well was added to Polysoap Immunoplate (Nunc). The recombinant protein was immobilized for 2 days at 4 °C. For serological examination of deer, plasma specimens frozen at −80 °C was diluted 1:100 with ImmunoBlock and used for ELISA. Horseradish peroxidase conjugated Protein G (Sigma) was used as 2ndaly antibody. OPD (o-Phenylenediamine dihydrochloride) tablets (sigma) was used as substrate for the detection of peroxidase activity. The absorbance at 450 nm was measured by a microplate reader (Multiskan FC, Thermo). OD value of rVenus (control) was subtracted as back ground. Mean OD values were derived from 3 independent experiments. For the ELISA positive threshold, the end-point cut-off was established by titration as the mean OD450 value of the plasma of PCR-negative deer, plus 3 standard deviations.

### Experimental animals

Specific pathogen free rabbits (JW strain) were purchased from Kitayama Rabes, Japan. Animal experimentation was carried out according to the Laboratory Animal Control Guidelines of National Institute of Infectious Diseases (institutional permission no. 116106).

## Results

### *Bd37* gene sequences of the isolates, IpSG13-13-, IpSG10 and MRNK

*Bd37* gene sequences were obtained by PCR cloning from genomic DNA (table 1) IpSG13-13-1, IpSG10 and MRNK and cDNA of IpSG13-13-1. Full length sequences were determined from 10 clones each (table 2) (total 40 clones). Accession numbers of representative sequences are shown in supplement table 2. Multiple alignments of the *Bd37* sequence showed heterogeneity and length polymorphism (supplement fig 1). Nevertheless, the various *Bd37* sequences from Japanese strains were distinctive and grouped by 3 types, namely JP-A, JP-B and JP-C types (table 2), among which sequence identities (%) were 69.89%-71.43% (supplement table 3). *Bd37* sequences of MRNK strain (EU-A type) were distinct from those of Japanese types. The identities between the JP types and EU-A type were 72-78% (supplement table 3). JP-A and EU-A were more polymorphic than others, and could be grouped by sub types, as JP-A1, A2 and A3 and EU-A1 and A2 respectively (table 2, supplement fig 1). Table 2 shows the number of plasmid clones carrying various sequence types. Genomic DNA and cDNA of IpSG13-13-1 contained JP-A and JP-B types, while genomic IpSG10 contained JP-A and JP-C types. European strain, MRNK, contained only EU-A (EU-A1 and EU-A2) type sequence.

### Comparison of Bd37 protein sequences

Putative protein sequences were obtained from all DNA sequences determined above. Percent identities (table 3) among the JP-A, JP-B and JP-C type protein sequences were distantly related (60.12-77.04%) to the reported sequences from *B. divergens* (Rouen1987, Y5, Weybridge and 71/07/B strains) and *B. capreoli*. The EU-A1 and A2 types from MRNK strain were closely related to the *B. divergens* Y5 strain (CAD48926) (89.91% and 99.69%, respectively) (table 3). All protein sequences determined in this study are shown in supplement fig 2A, 2B, 2C and 2D.

**Table 3.** Protein sequence percent identities.

| Isolates<sup>1</sup> | Accession numbers | Lineage<sup>2</sup> | Sequence type | 1 | 2 | 3 | 4 | 5 | 6 | 7 | 8 | 9 | 10 | 11 | 12 | 13 |
| --- | --- | --- | --- | --- | --- | --- | --- | --- | --- | --- | --- | --- | --- | --- | --- | --- |
| *B. divergens* IpSG13-13-1#5 | LC601771 | Asia lineage | JP-A1 | 100 | 98.19 | 92.17 | 63.22 | 64.42 | 70.72 | 69.11 | 70.72 | 74.92 | 75.23 | 76.76 | 59.45 | 61.45 |
| *B. divergens* IpSG13-13-1#10 | LC601780 | Asia lineage | JP-A2 | 98.19 | 100 | 93.67 | 63.22 | 64.72 | 70.4 | 69.11 | 70.4 | 74.92 | 75.23 | 76.76 | 59.15 | 61.45 |
| *B. divergens* IpSG10#2 | LC601804 | Asia lineage | JP-A3 | 92.17 | 93.67 | 100 | 64.13 | 64.42 | 69.47 | 67.89 | 69.47 | 74.62 | 74.92 | 77.68 | 59.15 | 60.24 |
| *B. divergens* IpSG13-13-1#26 | LC601787 | Asia lineage | JP-B | 63.22 | 63.22 | 64.13 | 100 | 60.54 | 65.44 | 66.67 | 65.44 | 58.98 | 59.28 | 59.21 | 52.73 | 52.08 |
| *B. divergens* IpSG10 #11 | LC601808 | Asia lineage | JP-C | 64.42 | 64.72 | 64.42 | 60.54 | 100 | 70.86 | 67.77 | 70.55 | 65.56 | 65.86 | 64.44 | 56.13 | 55.42 |
| *B. divergens* MRNK #5 | LC601796 | EU lineage | EU-A1 | 70.72 | 70.4 | 69.47 | 65.44 | 70.86 | 100 | 89.6 | 99.69 | 68.4 | 68.71 | 66.67 | 61.37 | 58.1 |
| *B. divergens* MRNK #7 | LC601790 | EU lineage | EU-A2 | 69.11 | 69.11 | 67.89 | 66.67 | 67.77 | 89.6 | 100 | 89.6 | 63.86 | 64.16 | 62.73 | 60.55 | 57.36 |
| *B. divergens* Y5 | CAD48926 | EU lineage |  | 70.72 | 70.4 | 69.47 | 65.44 | 70.55 | 99.69 | 89.6 | 100 | 68.4 | 68.71 | 66.67 | 61.06 | 57.8 |
| *B. divergens* Weybridge | CAD48924 | EU lineage |  | 74.92 | 74.92 | 74.62 | 58.98 | 65.56 | 68.4 | 63.86 | 68.4 | 100 | 99.71 | 87.92 | 54.88 | 54.55 |
| *B. divergens* Rouen1987 | CAD19563 | EU lineage |  | 75.23 | 75.23 | 74.92 | 59.28 | 65.86 | 68.71 | 64.16 | 68.71 | 99.71 | 100 | 88.22 | 55.18 | 54.84 |
| *B. divergens* 71/07/B | CAD48927 | EU lineage |  | 76.76 | 76.76 | 77.68 | 59.21 | 64.44 | 66.67 | 62.73 | 66.67 | 87.92 | 88.22 | 100 | 57.49 | 56.19 |
| *B. capreoli* 2770 | ACX30791 | B. capreoli lineage |  | 59.45 | 59.15 | 59.15 | 52.73 | 56.13 | 61.37 | 60.55 | 61.06 | 54.88 | 55.18 | 57.49 | 100 | 60.7 |
| *B. capreoli* 2801 | ACX30790 | B. capreoli lineage |  | 61.45 | 61.45 | 60.24 | 52.08 | 55.42 | 58.1 | 57.36 | 57.8 | 54.55 | 54.84 | 56.19 | 60.7 | 100 |

1. Genetic isolates are included
2. Lineage in the *B. divergens*/*B. capreoli* groupTable 4 Genetic diversity of *Bd37* JP-A, JP-B and JP-C types.

ClustalW alignments of representative protein sequences of JP-type and EU-type (Fig 1A) and JP-A1, A2 and A3 (Fig 1B) are shown in Fig 1A and 1B. Bd37 of Rouen87 strain (CAD19563) was included as reference (fig 1A). Regardless of the observed DNA sequence polymorphism, all putative protein sequences contained signal peptide at the N-terminus and the GPI anchor domain at the C-terminus with high probabilities (99.9%) (fig 1A, 1B). In other region, conserved and variable regions were mosaic throughout the sequences. High level of polymorphism was especially observed along forty amino acid residues after signal peptide sequence, where substitutions and in/dels were frequently occurred (fig 1A, supplement fig.1 and supplement fig.2). Cysteines involved in disulfide bridge and residues involved in salt bridge (Delbecq et al. 2008) were conserved among JP-A, JP-B, JP-C and EU-A with few exceptions (fig 1A and 1B).

**Fig. 1.**
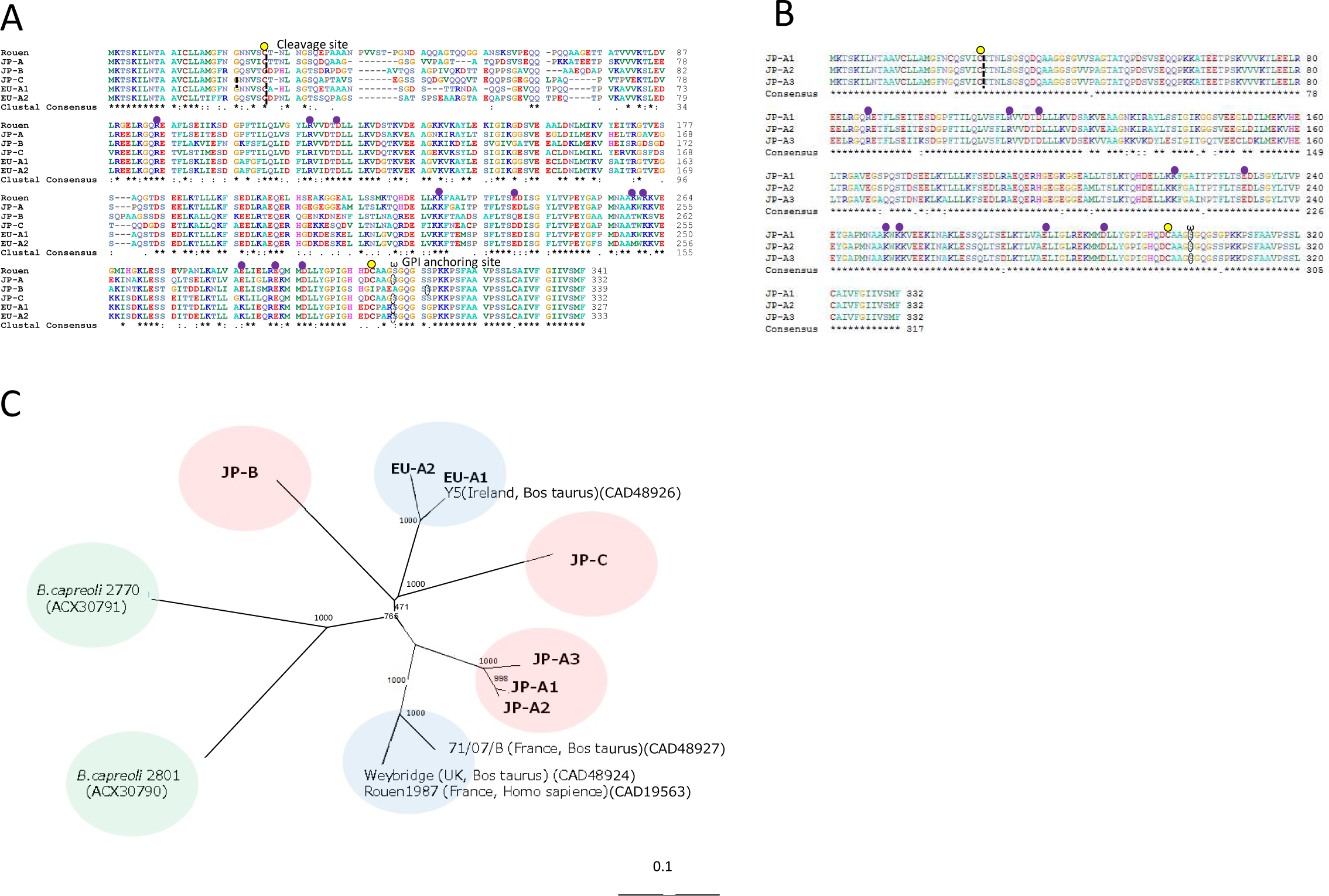
Alignment of Bd37 amino acid sequences (A, B) and phylogenetic tree (C). (A, B) Yellow and purple circles indicate disulfide bridge and salt bridge identified in Rouen1987 strain (Delbecq et al. 2002 and 2008). Cleavage site in signal peptide (solid line) and GPI anchoring site (circle) are shown (see method). (C) Phylogenetic tree of various Bd37 orthologs from the *Babesia divergens/B. capreoli* group. Boot strap values are shown in at each node. Sequences determined in this study are shown in bold. Red, blue and green circles indicate Bd37 from Asia lineage, EU lineage and *B. capreoli*, respectively.

Phylogenetic tree based on the clustalW alignment among the Bd37 proteins sequences determined in this study and from GenBank was shown in fig 1C. All Bd37 sequence types determined in this study (JP-A, JP-B, JP-C and EU-A) branched independently regardless of the lineages based on the *18S rRNA* (Zamoto-Niikura 2018). Bd37 from *B. capreoli* 2770 and 2801 strains (referred as Bcp37/41) were demonstrated to encode different proteins in size, 36.8kDa and 40.8kDa, respectively (Sun et al. 2011) and clearly separated from others (fig.1C).

The protein sequences of Bd37 were highly polymorphic (fig. 1A, 1B and supplement fig. 2) and independently evolved (Fig 1C). However, it seems not influence the propensity of the predicted liner B cell epitope probability (Fig 2A). The pattern was conserved throughout the sequences of all Bd37, except for those of *B. capreoli* 2801. By using the default cut-off value of 0.5 as suggested in the program, the propensity for B cell epitopes was abundant and encompasses almost all of the entire protein, except for N- and C-terminal parts (fig 2A). Regions where probability was relatively high (score >0.6) were indicated as epitope A, B and C (fig 2A). Sequence alignments of the epitopes, except for that of *B. capreoli* 2801, were shown in fig 2B. Protein sequence in epitope A were more polymorphic than those in epitope B and epitope C.

**Fig. 2.**
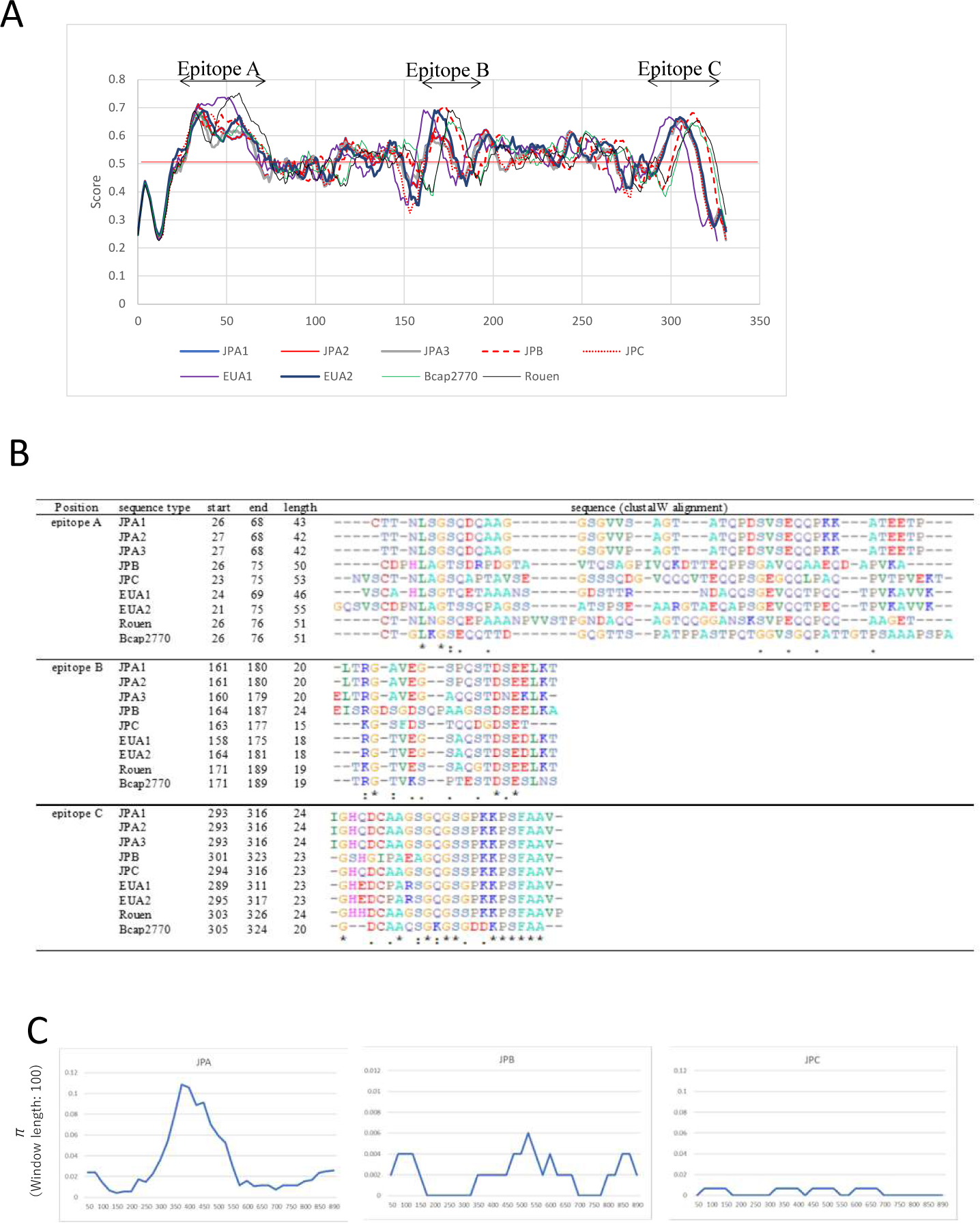
B-cell epitopes of various Bd37 orthologs predicted by BepiPred-2.0 server and window graph of DNA polymorphism in JP-A, JP-B and JP-C. (A) The threshold (default value =0.5) is shown as red line. Predicted B cell epitope above 0.6 are designated as Epitope A, B and C. Sequence types with accession numbers are shown in table 3. (B) Amino acid sequences of the Epitope A, B and C. (C) Nucleotide diversity of JP-A, JP-B and JP-C examined by sliding window plot with a window length of 10bp and a step size of 25bp. Accession numbers of nucleotide sequence used are shown in table4.

**Fig. 3.**
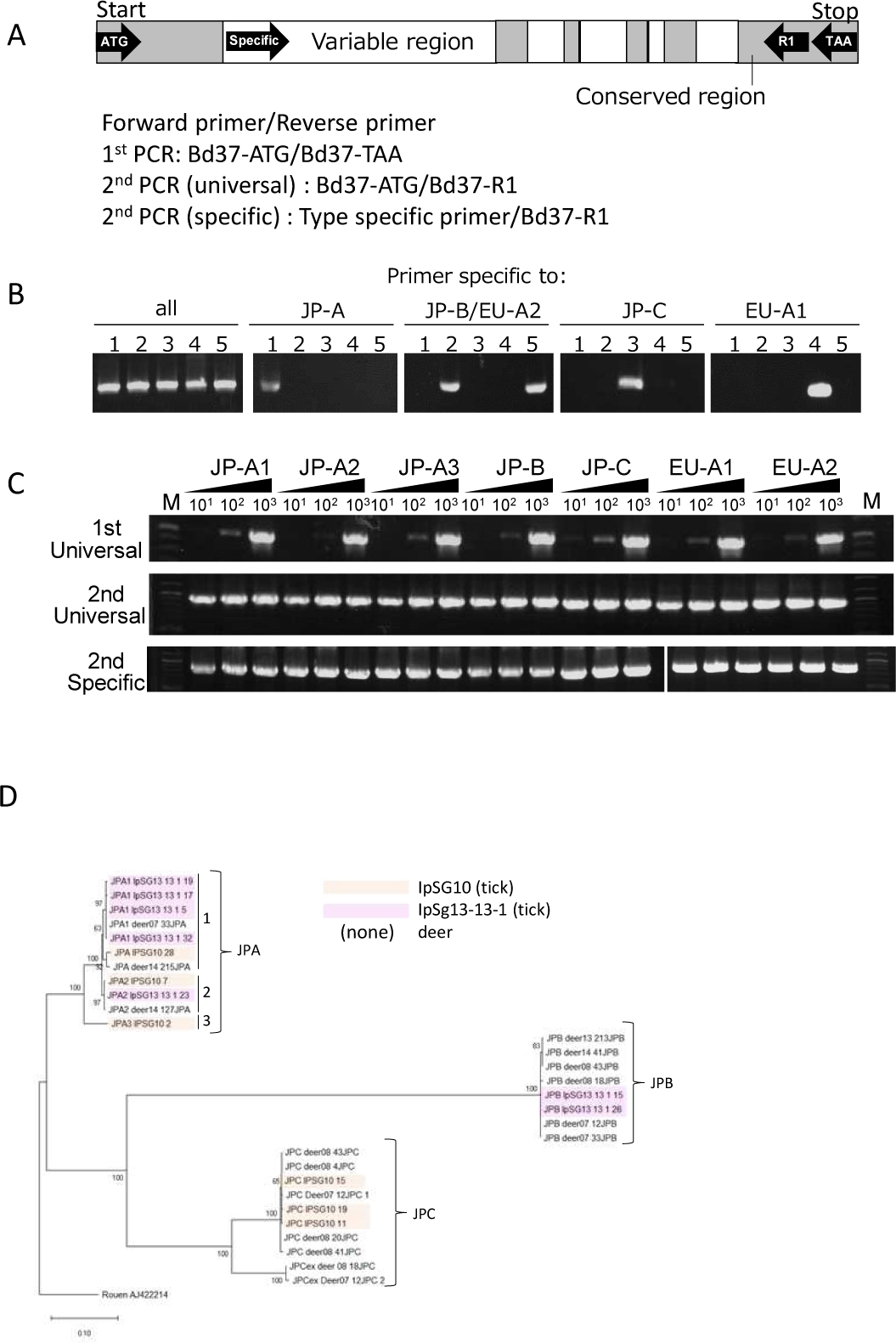
Type-specific PCR targeting the Bd37 genes of *B. divergens* Asia and EU lineage. (A) Schematic diagram of the PCR primer design. Black arrows show primers for primary and second PCR Black and white boxes indicate conserved and variable regions. Primer sequences are shown in supplement table S1. (B) Specificity of primers. (C) Specificity of the type-specific PCR. PCR amplification using universal primers Bd37-ATG, or specific primers Bd37-JPA F1, Bd37-JPB/EUA2 F2, Bd37-JPC F1, or Bd37-EUA1 F1) and Bd37-TAA. Plasmids carrying the each Bd37 gene of genetic strains shown in were used as the template. M, marker. (C) Sensitivity of specific nested PCR. Conventional PCR using universal primers Bd37-ATG and Bd37-TAA or Bd37-R1 (upper and middle panel, respectively) or s Bd37-JPA F1, Bd37-JPB/EUA2 F2, Bd37-JPC F1, or Bd37-EUA1 F1) and Bd37-TAA (lower panel) with the plasmid carrying the Bd37 gene described above. Plasmids were diluted 10-fold from 1×10^1^ to 1×10^3^ copies and used (lanes 10^1^ to 10^3^). M, marker. (D) Phylogenetic trees of Bd37 JP-A, JP-B and JP-C DNA and protein sequence from sika deer and *I. persulcatus* tick. Boot strap values are shown in at each node.

### Genetic analysis

Since protein sequences among JP types were highly polymorphic, we further examined segregating polymorphism among gene sequences (table 4 and fig 2C). Note that only 3 of JP-C type DNA sequence were available. By comparison of JP-A and JP-B, haplotype diversity was comparable (0.897±0.056 vs 0.933±0.077), while number of segregating sites (S) (74 vs 9) and nucleotide diversity (Pi) (0.0279 vs 0.0019) were relatively higher in JP-A. It indicates that multiple nucleotide substitutions within a single JP- A gene sequence frequently occurred and possessed higher levels of genetic diversities. The ratio of non-synonymous (dN) to synonymous (dS) substitutions (dN/dS) was calculated to asses natural selection acting on Bd37. The ratio of JP-A was dN/dS <1, representing potential evidence that a purifying selection pressure might be taking place to shape the populations. This negative (purifying) selection was statistically significant examined by Z-test (dN<dS, P= 0.00097). dN/dS value of JP-B was slightly over of 1 (1.044), but neither of positive (dN>dS) nor neutral (dN=dS) selections was statistically significant. Fig. 2C shows sliding window plot with a window length of 10bp and a step size of 25bp providing detailed analysis of the diversity. Pi of JP-A was range from 0.0429 to 0.19, and the highest peak diversity was observed within the nucleotide position at 300-500 (actual position at 335-535 of CDS). The highly divergent area, 335-535 of CDS, is correspond to position at 112-145 of proteins sequence.

**Table 4.** Genetic diversity of Bd37 JP-A, JP-B and JP-C types.

|  | no | length | popsition | h | hd | SD | S | Pi | dN/dS | p (Z-test) |
| --- | --- | --- | --- | --- | --- | --- | --- | --- | --- | --- |
| JPA | 17 | 930 | 34-963 | 10 | 0.897 | 0.056 | 74 | 0.02786 | 0.334 | 0.00097* dN<dS |
| JPB | 10 | 951 | 34-984 | 8 | 0.933 | 0.077 | 9 | 0.00189 | 1.044 | 0.31910 dN>dS |
| JPC | 3 | 930 | 34-963 | 3 | 1 | 0.272 | 4 | 0.00287 | 0.295 | 0.21027 dN<dS |

Number of Haplotypes, h; Haplotype (gene) diversity, Hd; Number of polymorphic (segregating) sites, S; Nucleotide diversity, Pi; Total number of synonymous changes, dS; Total number of nonsynonymous changes, dN

### Development of discriminative PCR

For detailed molecular analysis of *B. divergens* in deer, a discriminative PCR was developed (Fig 3A) by designing primers specific for variable regions in JP and EU types (supplement table 1). The specificity of this PCR system was examined by using plasmids encoding each type. The results shown in Fig 3B, demonstrating formation of a single positive signal only when the specific primer matched, except for JP-B/EU-A2. When the JP- B/EU-A2 primer was used, amplicons were generated from the plasmids encoding JP-B and EU-A2. Thus, we confirmed the sequence type, JP-B or EU-A2< by direct sequencing. Sensitivity of the type-specific PCR was examined by using 10-fold dilutions of plasmid (1×10^1^ to 1×10^3^ copies in a PCR mixture) as template in the nested PCR. In the first-round PCR (Fig 3C top panel) by using universal (top panel) primers, products were observed when 1×10^2^ and 1×10^3^ copies were used as template. In the nested PCR, in which the first-round PCR products of 1×10^1^ to 1×10^3^ copies respectively were used as template, amplicons were visible at all concentrations (Fig 3C, middle and bottom panels) regardless of using universal or specific primers.

### Examination of the field samples

The discriminative PCR developed above was applied to examine *B. divergens* in wild sika deer (*Cervus nippon,* erythrocytes) (22 samples) and additional field collected tick (14 samples) (table 5). The samples of deer and ticks were previously tested for presence of *B. divergens* by 18S rRNA based PCR (Zamoto-Niikura 2014, 2018, 2020). By using the primers targeting all *Bd37* sequence types (universal), PCR amplicons were indeed developed from all samples. Further discriminative PCR revealed JP-A specific amplicons were exclusively identified in all 36 samples, while JP-B/EU-A2 and JP-C specific amplicons were generated only from 17 samples, respectively. EU-A1 specific amplicon was not detected. Table 5 shows the number of positives in each combination pairs.

**Table 5.** Summary of field survey examined by discriminative PCR.

| Species | Materials | No. PCR positive |  |  |  | Total |
| --- | --- | --- | --- | --- | --- | --- |
|  |  | A only | A and B | A and C | A, B and C |  |
| *Ixodes persulcatus* | Whole body | 2 | 3 | 5 | 2 | 12 |
|  | Salivary glands | 1 | 1 | 0 | 0 | 2 |
| *Cervus nippon* | Erythrocytes | 5 | 6 | 5 | 6 | 22 |
| Total |  | 8 | 10 | 10 | 8 | 36 |

The amplicons amplified from deer sample by the discriminative PCR above were further sequenced to examine single nucleotide polymorphisms (SNPs) within individual genomic samples (supplement fig 3A, 3B and 3C). Among 12, 7 and 6 of JP-A, JP-B and JP-C, respectively, SNPs were observed in 8 and 1 of JP-A (n=8) and JP-C (n=1) (Supplement fig 3A, 3C). SNP was not observed in the JP-B (supplement fig 3B). Length polymorphism was observed only in a JP-C (deer 08#18) where the length is 9nt shorter than others (supplement fig 3C). Furthermore, JP-C from deer 07#12 contained high polymorphic region in latter half of the sequence (87/335) (supplement fig 3C). Further examination of 07#12 sequences by cloning revealed that (Supplement fig 3D), 2 sequence types were mixed (JPC-1 and JPC-2) (supplement fig 3D). Identities between the two deduced amino acid sequences was 80-84%.

By using all haplotype sequences from deer and ticks in this study, phylogenetic tree (DNA, 738nt) was constructed (fig 3D), demonstrating the genetically closely related parasites were transmitted between the tick vector *I. persulcatus* and reservoir *Cervus nippon*. The presence of sub lineages within JP-A and JP-C clades may indicate growing population and genetic drift.

### Expression of recombinant Bd37 proteins

To examine whether Bd37 was antigenically recognized in deer, recombinant Bd37 proteins (rBd37) fused with thioredxin (Trx) at N-terminal were expressed in *E. coli*. The rBd37 of all types and control (rVenus) were expressed as soluble proteins and purified under native condition. The SDS-PAGE and western blot analysis using anti-Trx antibody (fig 4A) showed that all recombinant proteins were expressed as expected size. The observed proteins were 11.7 KDa larger to the expected size because of fused thioredoxin.

**Fig. 4.**
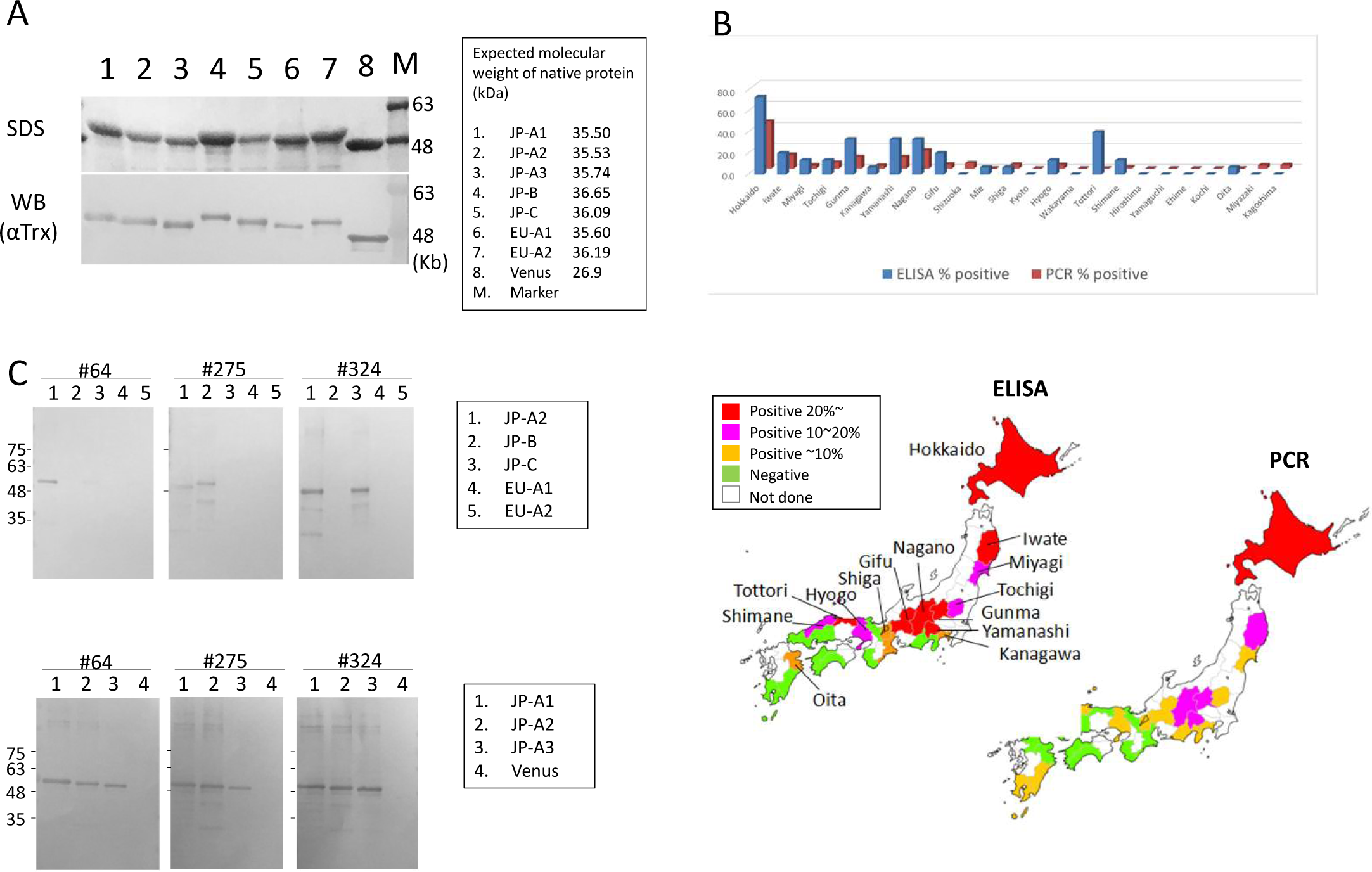
Expression, cross-reactivity and use in ELISA of recombinant Bd37 from *B. divergens* Asia and EU lineages. (A) SDS PAGE and western blot analysis of purified rBd37 proteins conjugated with Thioredxin (Trx) and His-tag. Note that molecular weight of Trx and His-tag is 16KDa. Anti Trx antibody was used as primary. (B) Wild sika deer were serologically examined by rJP-A based ELISA (n=360). Positive rates in each prefecture (n=24) are shown. PCR positive rate is shown for comparison. Positive rates shown by bar chart and geographical map (upper and lower panel, respectively) (C) Different serological profile of deer (upper panel) and cross-reactivity of rBd37 (lower panel). Number above the line indicates designated sample number of wild sika deer. Protein A/G was used for 2nd antibody.

### Establish ELISA system based on rBd37 JP-A

Genetic analysis revealed JP-A1 and JP-A2 were exclusively detected in all *B. divergens* examined in Japan (table 2, fig 3D). Thus, JP-A2 was used as representatives of rBd37 JP-A as antigen in the ELISA. Rabbit serum raised against rBd37 JP-A2 was cross reactive to all JP-A types examined by western blot and ELISA (data not shown).

The presence of cross-reactive antibody in deer against the rBd37 JP-A was examined by using plasma from PCR positive (50 specimens) and PCR negative deer (12 specimens), latter of which were hunted in *B. divergens*-PCR free prefectures (Zamoto-Niikura 2020). In the ELISA, mean OD values of PCR positive and negative deer were 0.12±0.16 and 0.002±0.01, respectively. Out of 50 PCR positive deer examined, OD values of 32 samples were above the end-point cut-off value, which was established by the mean OD value of the PCR negative plasma plus 3 standard deviations (mean + 3SD).

### Serological screening of *B. divergens* infection in deer

By using the rBd37 JP-A based ELISA, antibody in wild sika deer in Japan were examined. Fifteen deer plasma each were randomly chosen from 24 prefectures (total 360). Overall positive rate of the serological test was 13.9%, which was approximately twice as high as the positive rate derived from PCR tests (6.8%) (Zamoto-Niikura 2014, 2020). Higher positive rate (>20%) (fig 4B) were observed in Hokkaido and prefectures having mountain area including Iwate, Miyagi, Gunma, Yamanashi, Nagano and Gifu. The trend of the positive rate among prefecture was comparable to that examined by PCR (fig 4B) (Zamoto-Niikura 2020).

### Detection of antibodies against rBd37 JP-B and JP-C in deer

Addition to antibody against rBd37 JP-A, those of rBd37 JP-B and JP-C in deer were examined by western blot analysis. Three deer plasma showing relatively high OD value (#64, #275 and #324) in the JP-A based ELISA were tested (fig 4C). Major bands were recognized at approximately 50kDa of rBd37 JP-A in all specimens tested. Additional bands against rBd37 JP-B and JP-C were also recognized in the plasma of #275 and #324, respectively. Recombinant Bd37 of JP-A2 as well as rBd37 JP-A1 and A3 also reacted to all plasma specimens of wild sika deer, suggesting JP-A1, A2, and A3 could be considered antigenically similar in this system.

## Discussion

To seek immunodiagnostic gene and antigen of Asia lineage in the *Babesia divergens/B. capreoli* group, we investigated GPI anchored protein Bd37, a major surface protein of *B. divergens* sensu strict. Direct examination of parasite’s genome in the vector (table 1) and reservoir (table 5) identified unique and novel features of the Bd37 gene family: 1. Unlike other lineage, the Asia lineage carries as much as three distinct Bd37 ortholog genes, namely *Bd37 JP-A, JP-B* and *JP-C* (paralogs) (table 2). 2. Genetic diversities of them were distinct (table 4) but B cell epitope was similar (fig 2). 3. The *JP-A* was exclusively encoded in all genomes isolated, while *JP-B* and *JP-C* were encoded 50% of the genomes examined, respectively (table 2and 5). 4. Antibodies against recombinant JP-A (rJP-A), JP-B and JP-C were evident in sika deer indicating expression of all Bd37 orthologs in blood stage (fig 4A). 5. Although the sequences were highly polymorphic, the purified rJP-A2 was widely cross reactive to the JP-A variants (fig 4C). Finally, we regarded JP-A as diagnostic marker and established rJP-A2 based ELISA (fig 4B). This system demonstrated that over all positive rate of 13.9% in wild sika deer throughout Japan (24 prefectures, n=360) was 2 times higher than that examined by PCR.

*B. divergens* parasites have been isolated from mammals including human and bovine, and maintained erythrocytes in vitro or laboratory animals. Such efforts established many strains and studies for Bd37 performed on them (Gorenflot 1990, Delbecq 2002, Hadj-Kaddour 2007, Sun 2011, Gonzalez 2015, Bastian 2012, Gabrielli 2012, Hall 1999, Rozas 2017, Kumar 2018, Delbecq 2008). Unfortunately, the Asia lineage has not biologically isolated. Thus, in primary investigation, we utilized salivary grands of *Ixodes persulcatus* ticks as gnomic resource and isolated as much as three type genes (JP-A, JP-B and JP-C) from only two ticks (table 2). Further investigation of deer (table 5) did not identify additional sequence type. In the salivary glands, multinucleated sporoblast matures and multiple sporozoites develop. Based on the calculation from semi-quantitative PCR (fig. 3C), original salivary grands (whole) contained 5×10^5^< copies /tick (data not shown). This high-number of parasite’s population were rich in diversity and advantageous for deep genomic analysis. Furthermore, isolation of mRNA of JP-A and JP-B from the salivary glands (table 4) indicates expression of JP-A and JP-C in the sporozoite stage and need for erythrocytes infection in next stage (due to for lack of samples, isolation of JP-C mRNA could not be attempted). Although it is quite bothersome to obtain salivary glands infected with matured, multiplied *Babesia* parasite (activated), this study showed the salivary glands as informative tool to investigate not only interaction between vector ticks and mammals but also population genetics.

Genetic polymorphisms are nature of genes encoding merozoite surface proteins. The polymorphism of *B. divergens Bd37* gene was primary reported by Hadj-Kaddour et. al (Hadj-Kaddour et al. 2007), where fourteen isolates of *B. divergens* were clearly distinguished by five major polymorphic groups examined by PCR-Restriction fragment length polymorphism (RFLP) using *Rsa*I and *Bgl* II restriction enzymes. Sun et al. (Sun et al. 2011) performed the analysis similarly and observed extreme polymorphism within *B. capreoli,* which was greater than those among *B. divergens* isolates. In our study, JP type sequences indeed included the *Rsa*I and *Bgl* II recognition sites which were unique to JP-A, JP-B, JP-C, as well as EU-A (supplement fig 1). However, *Rsa*I recognition site is absent in some JP-A PCR clones suggesting use of the PCR-RFLP in Asia lineage may give us incomprehensive result. PCR-RFLP is fast and simple method for typing and widely used.

While it should be carefully used in sample directly prepared from wild animal (Thompson et al. 2022). PCR cloning followed by sequencing (or direct sequencing) as shown in this study should be ideal for detail investigation of genetic polymorphism and identification of the polymorphic groups (referred as “type” in this study).

This study was the first ecological survey of using recombinant surface protein of Piroplasm as diagnostic antigens in wildlife, deer (fig 4). Research and management of wildlife diseases are recognized as a part of controlling One Health. One of most important wild life is Cervid, which has increased its population worldwide (Abrantes and Vieira-Pinto, 2023), and infected with various genotypes of zoonotic (or zoonotic potential) microorganisms. *Toxoplasma gondii*, a globally distributed zoonotic protozoa, infects white-tailed deer (*Odocoileus virginianus*) with three genotypes/haplogroups including a human pathogenic type in the USA (Dubey et al. 2020). *Anaplasma phagocytophilum*, a tick-borne pathogen of granulocytic anaplasmosis (HGA) in humans, dogs and horses, infects roe deer (*Capreolus capreolus*), red deer (*Cervus elaphus*) and Iberian red deer (*Ce. Elephus hispanicus*) with various genotype in cluster I(*ankA*) and Ecotype I(*groEL*) in Europe (Matei et al. 2019). Piroplasm had thought to strictly infect specific host species. However recent studies suggest it jumps species barrier including cervid and human. *Babesia* spp. in patients in China (Wang 2019) carry18S rRNA sequences (KX839233 and MK256977, respectively) identical to those of *B. odocoilei* (ex. FJ944828), a very common piroplasm of cervid.

Identical *B. odocoilei* 18S rRNA was also found in horse in Italy (Zanet 2017). In Japan, parasite closely related to *B. odocoilei,* another piroplasm of cervid, infects dogs (Inokuma et al. 2003, Kubo et al. 2015, Yamasaki et al. 2021). The Asia lineage studied in this study is primary transmitted by *I. persulcatus* (Zamoto-Niikura et al. 2014, 2018), which is one of the important tick species in Eurasia in terms of public health. No human cases of the Asia lineage infection have been found so far, a potential risk of infection in human can’t be ignored. Continuous investigation of the Asia lineage in sika deer, as well as ruminants and companion animals may help to detect any threats from potential of zoonotic transmission to human.

In the analysis of genetic diversity (table 5) some characteristic features were observed in JP-A. 1. higher values of nucleotide diversity (π). 2. High number of segregating sites. 3. Significantly low dN/dS value (0.334). These results indicate that nucleotide substitutions occurred frequently without significant structural change. Indeed, cross reactivity among JP-A1, JP-A2 and JP-A3 was observed in the western blot (fig 4C) and ELISA (data not shown). Sliding window plot (fig 2C) showed nucleotide diversity was increased from 300 to 500 which is corresponding to the amino acid residues at 100 to 166 (fig 1A,1B and 2A). We assume that JP-A possess strong selective pressure due to a possible role of RBC adhesion (Delbecq 2008). While, genetic polymorphism should be assured as major surface protein to escape host immunity. We suppose combination of different properties, invasion and escape (Cuy-Chaparro et al. 2022) have shaped the unique Bd37 JP-A genetic polymorphism under different selective pressures.

In conclusion, Bd37 exhibits extensive sequence diversity in Asia lineage. Among 3 types identified, one was stably and others were sporadically retained in the genome.

Although some specific futures of Bd37 are conserved, dN/dS value and nucleotide diversity of JP-A was significantly different from those of JP-B and JP-C indicating mechanism of intragenic recombination and natural selection may be different. Recombinant JP-A based ELISA was sensitive and ideal for environmental survey. *B. divergens* group includes multiple lineage zoonotic and zoonotic potential. Continued surveillance and research of *B. divergens* group and its effects on cervid ecosystems is vital for controlling the long-term consequences of this emerging disease.

## Acknowledgments

Special gratitude is extended to Dr. Chiaki Ishihara for his continuous support and thoughtful guidance throughout this study. This work was supported by AMED under Grant Number JP 21fk0108097j0803, 20fk0108097j0802, 19fk0108097j0201 and JSPS KAKENHI Grant number 18K06398.

