## Supplemental for "Genetic investigation of GPI anchored Bd37 orthologs in *Babesia divergens* group and use of recombinant protein for ecological survey in deer"

### Supplement table S1-3

Table S1. Primers used in this study

| Name | Forward/reverse | Sequence (5' to 3') | Specificity |
|---|---|---|---|
| Bd37-ATG | forward | ATGAAAACCAGTAAGATTCTCAACACTGCTGCT | universal |
| Bd37-TAA | reverse | TTAGAACATTGAYACAATGATACCGAASACAATGGC | universal |
| Bd37-TAA removed | reverse | GAACATTGAYACAATGATACCGAASACAATGG | universal |
| Bd37-SQF1 | forward | CCTTCGTGTTRTYGACACAGATCTTCTCCTG | universal |
| Bd37-JPA F1 | forward | CAATCTGTAATCTGCACCACCAATCTYAGCGGTTC | JP-A specific |
| Bd37-JPB/EUA2 F2 | forward | TTACCTGTGATCCACATCTCGCCGGCACAAGC | JP-B, EU-A2 specific |
| Bd37-JPC F1 | forward | GCAGGCTCACAGGCGCCAACAGCAGTTAGTGAAGG | JP-C specific |
| Bd37-EUA1 F1 | forward | CAGGTGATTCAACTACTAGGAATGATGCGCAGCAGT | EU-A1 specific |
| Bd37-R1 | reverse | GCGAAGGATGGCTTCTTMGGACYAGATCCCTG | universal |

Table S2. Accession numbers

| Sequence type | Strain | CDS length | Accession number |
|---|---|---|---|
| JP-A1 | IpSG13-13-1 | 999 | LC601771 |
| JP-A2 | IpSG13-13-1 | 999 | LC601780 |
| JP-A3 | IpSG10 | 999 | LC601804 |
| JP-B | IpSG13-13-1 | 1020 | LC601787 |
| JP-C | IpSG10 | 999 | LC601808 |
| EU-A1 | MRNK | 984 | LC601790 |
| EU-A2 | MRNK | 1002 | LC601796 |

Table S3. Percent identity matrix (DNA)

| | Sequence type | Accession numbers | 1 | 2 | 3 | 4 | 5 | 6 | 7 |
|---|---|---|---|---|---|---|---|---|---|
| 1 | JP-B | LC601787 | 100.00 | 70.00 | 70.00 | 69.89 | 71.40 | 72.02 | 74.60 |
| 2 | JP-A3 | LC601804 | 70.00 | 100.00 | 94.09 | 94.52 | 71.43 | 72.72 | 71.52 |
| 3 | JP-A1 | LC601771 | 70.00 | 94.09 | 100.00 | 98.82 | 71.32 | 74.59 | 72.92 |
| 4 | JP-A2 | LC601780 | 69.89 | 94.52 | 98.82 | 100.00 | 71.21 | 74.59 | 73.03 |
| 5 | JP-C | LC601808 | 71.40 | 71.43 | 71.32 | 71.21 | 100.00 | 77.85 | 75.62 |
| 6 | EU-A1 | LC601790 | 72.02 | 72.72 | 74.59 | 74.59 | 77.85 | 100.00 | 92.35 |
| 7 | EU-A2 | LC601796 | 74.60 | 71.52 | 72.92 | 73.03 | 75.62 | 92.35 | 100.00 |

### Supplement fig. 1

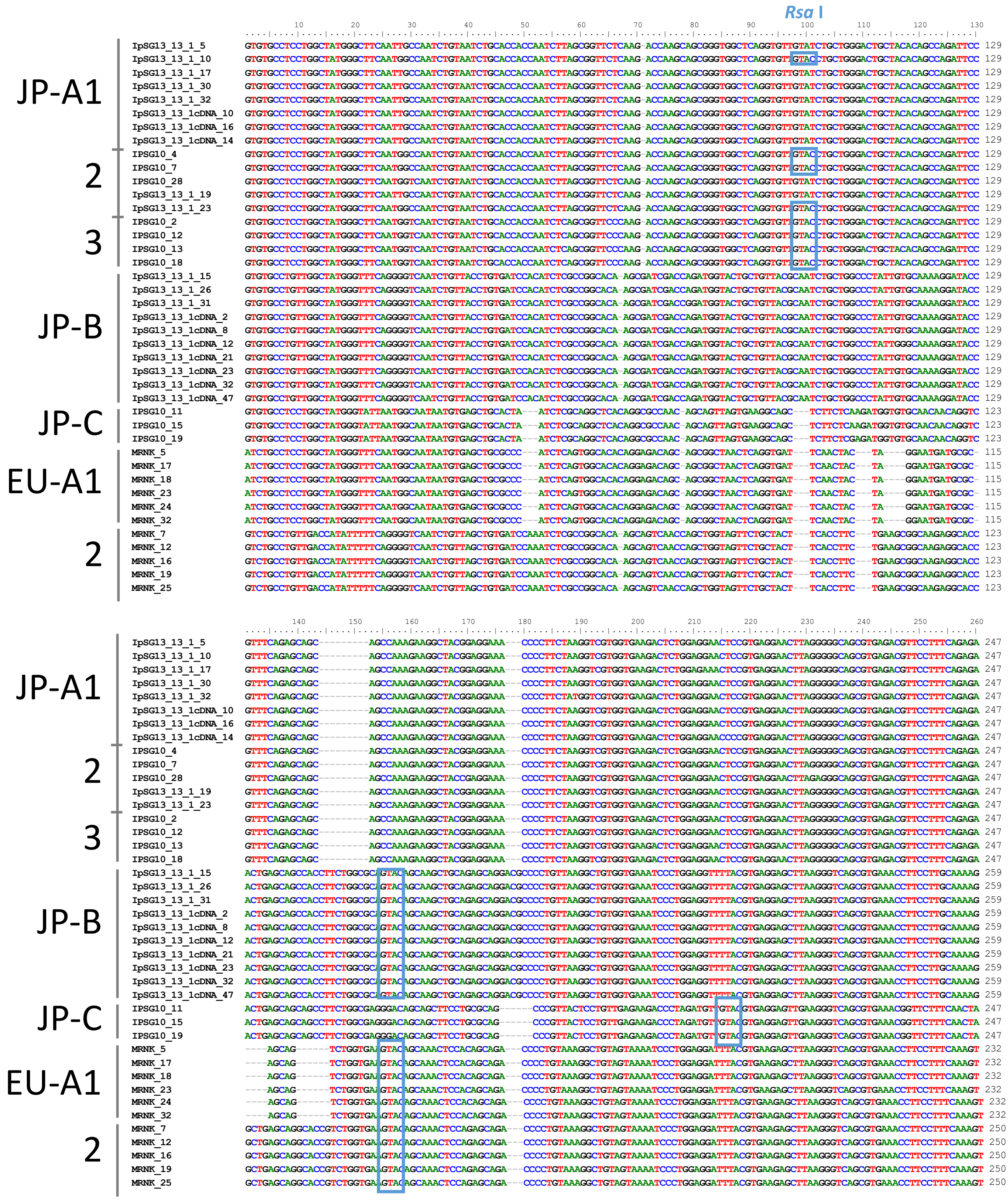

Supplement fig. 1 cont.

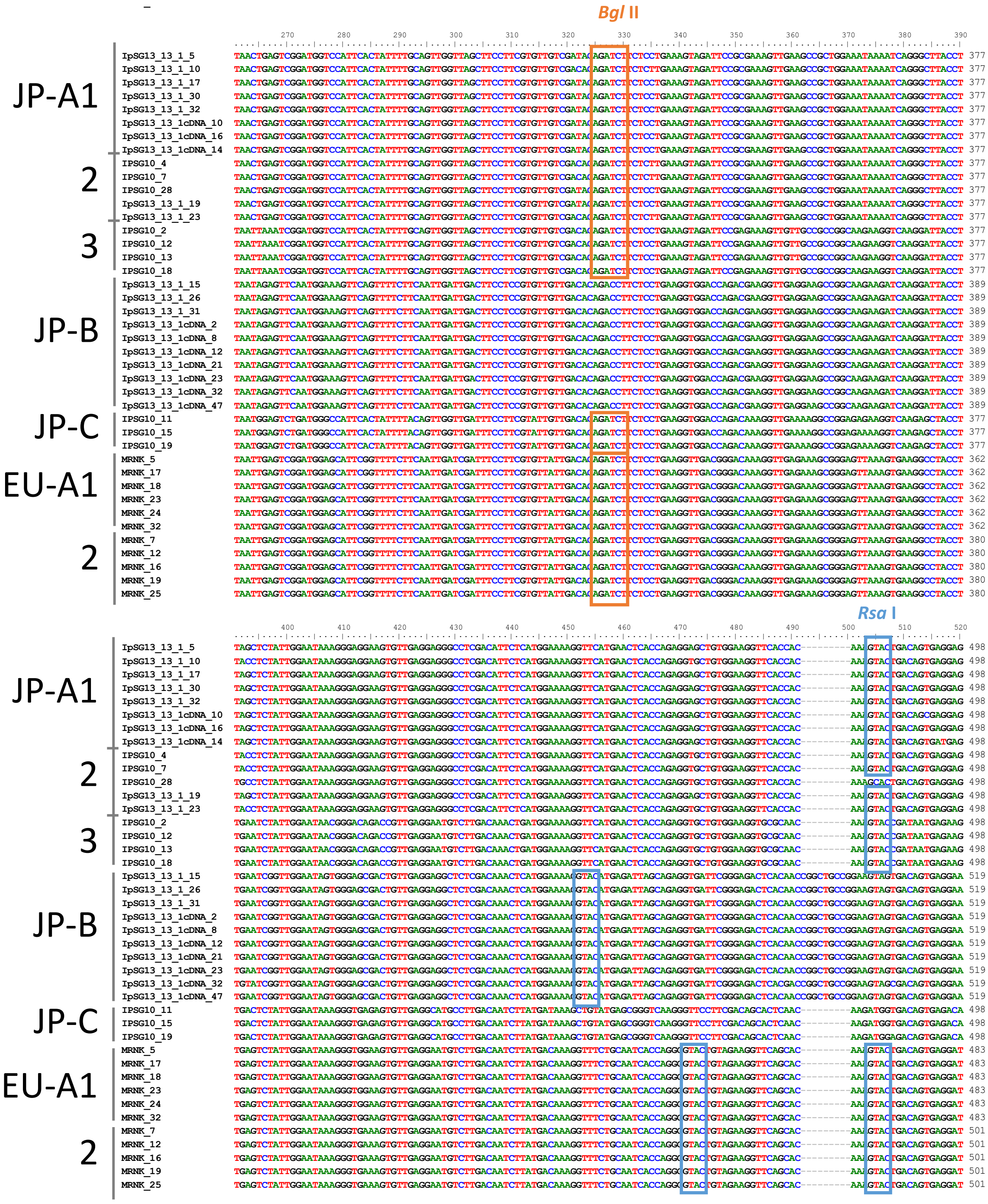

### Supplement fig. 1 cont.

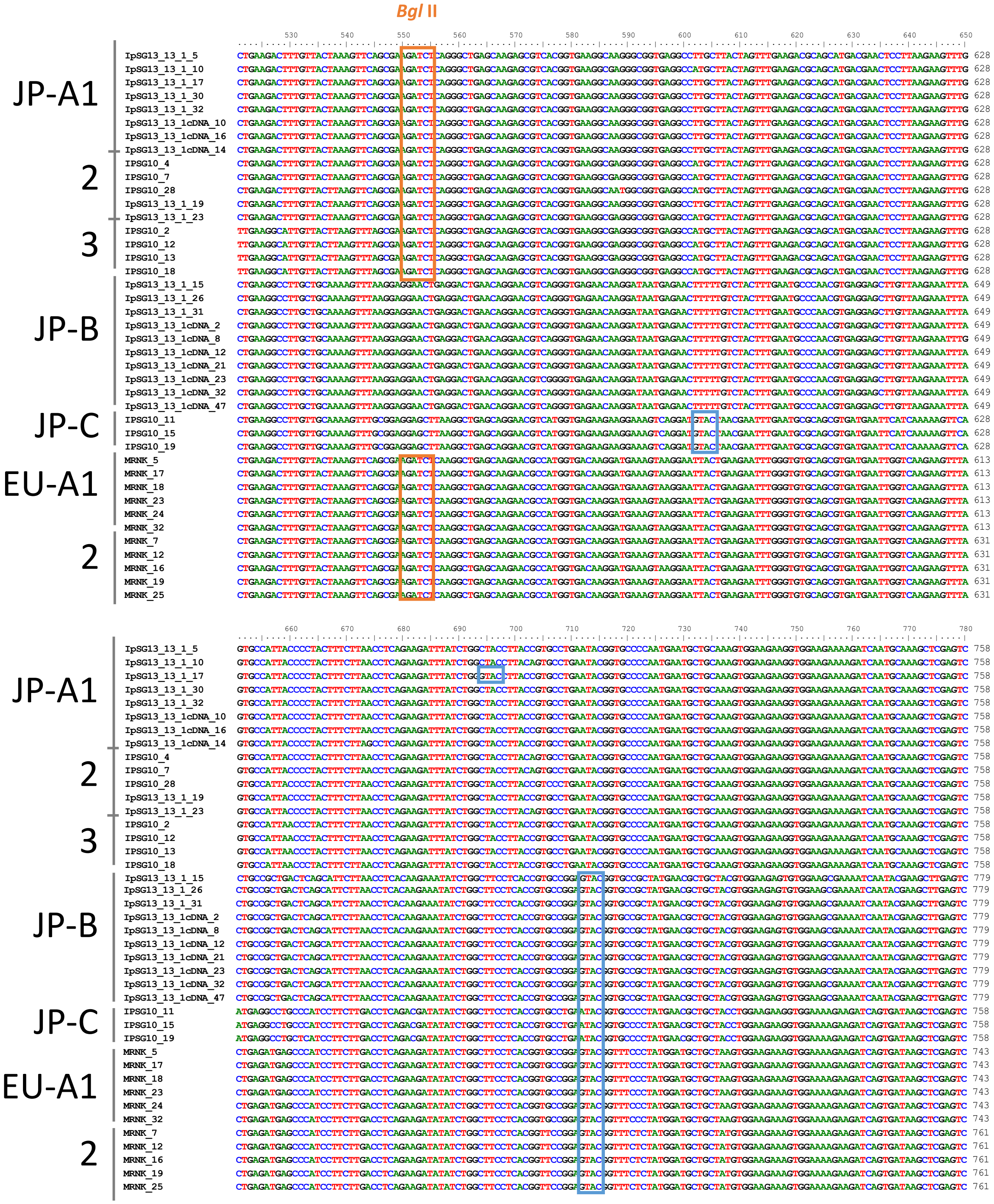

Supplement fig. 1 cont.

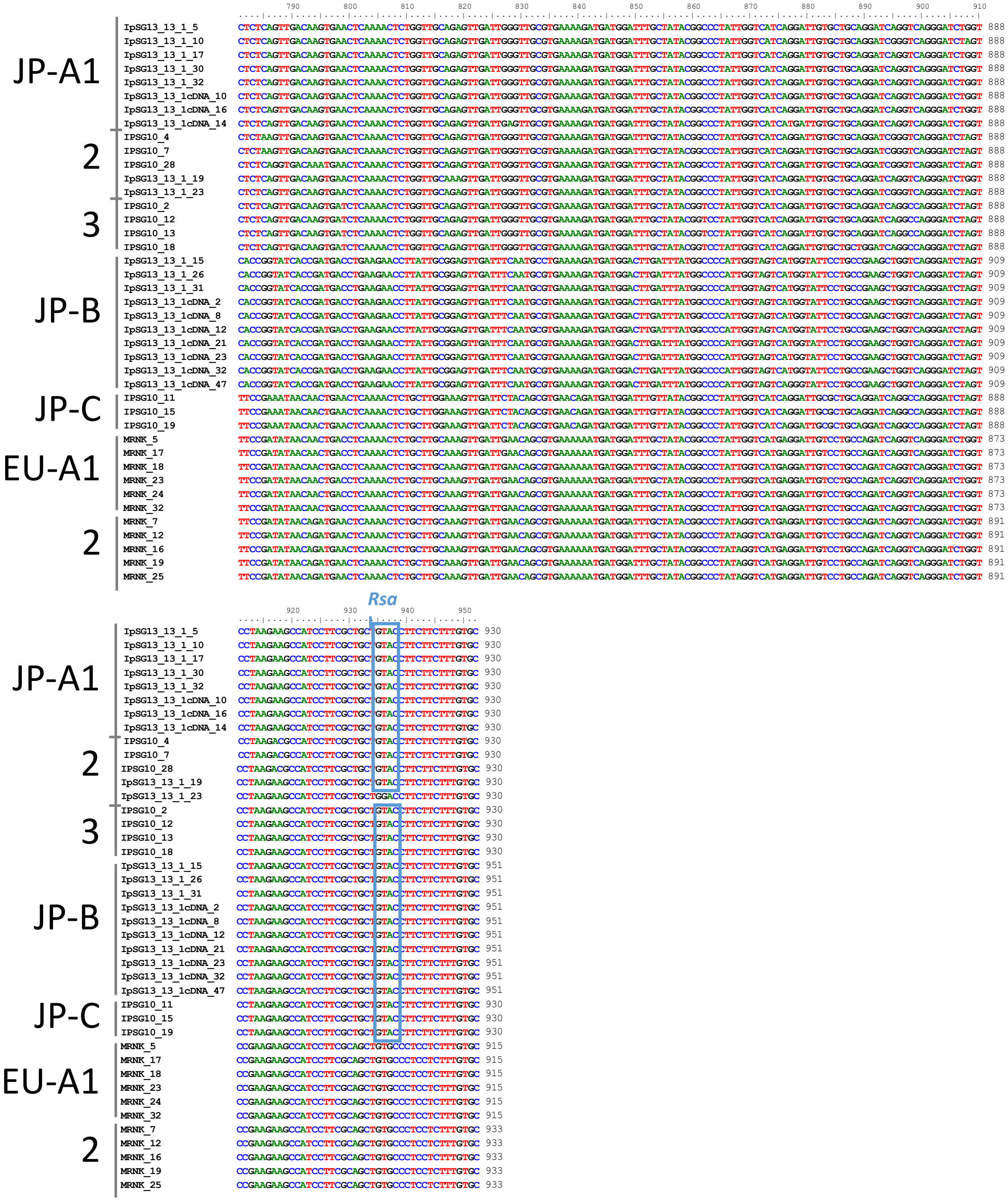

Supplement fig. 2A

JP-A protein sequence

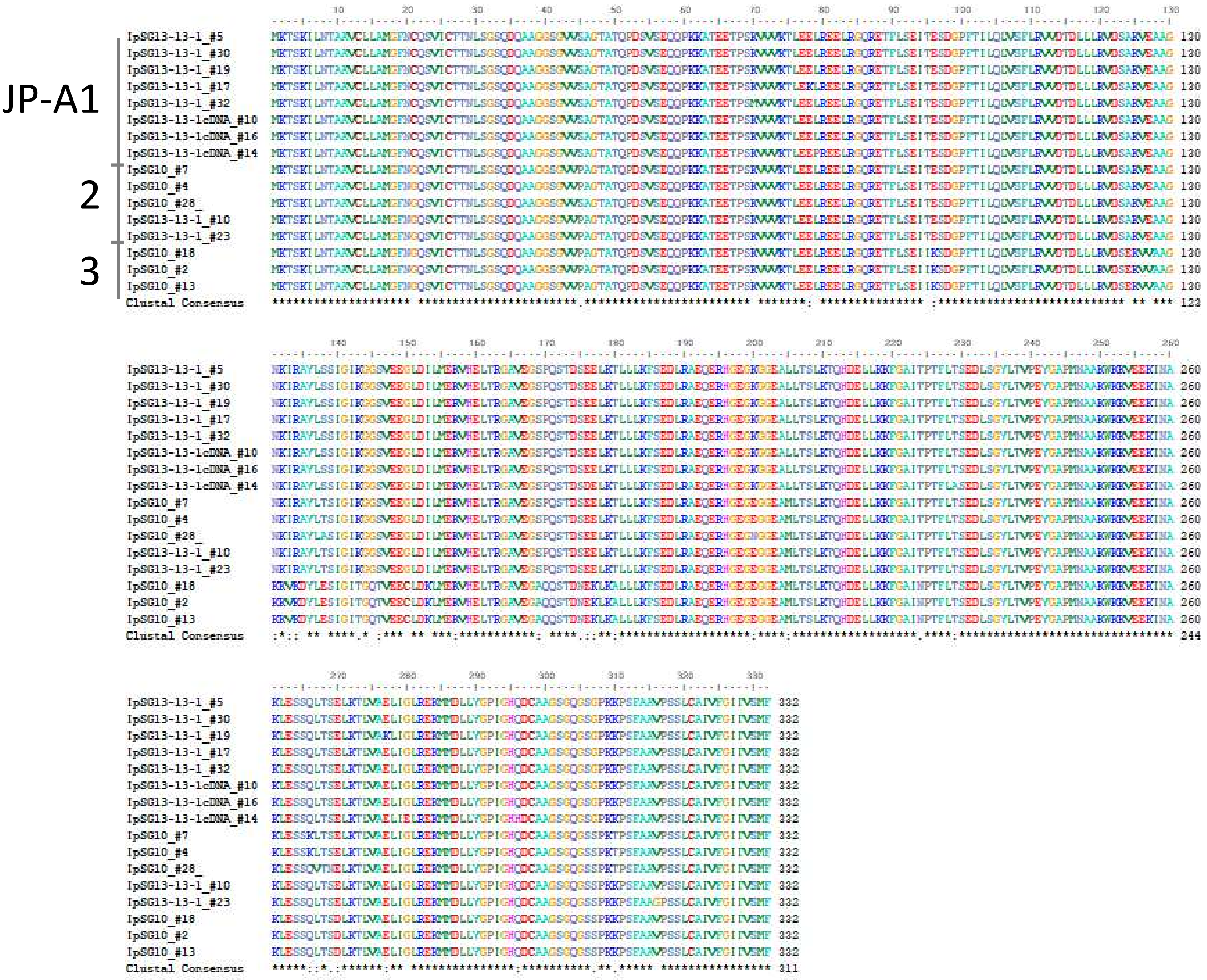

Supplement fig. 2B

JP-B protein sequence

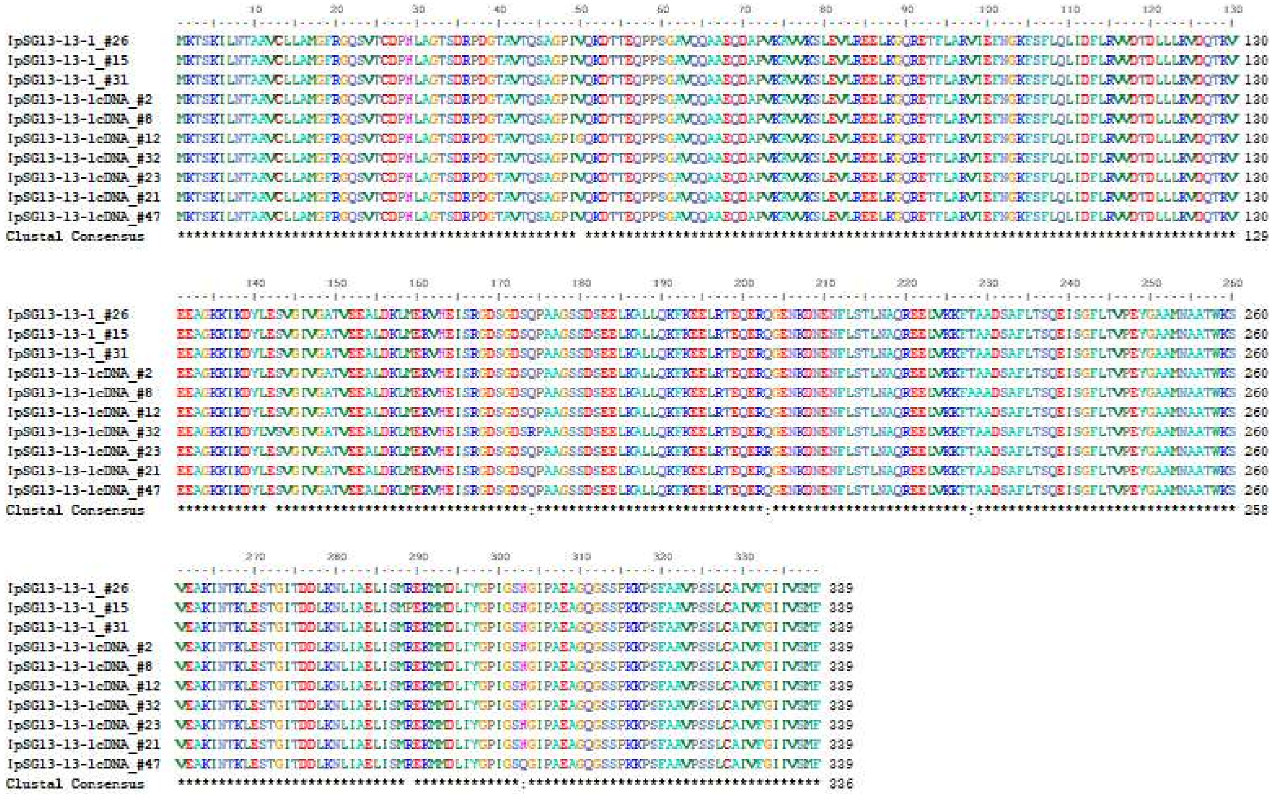

Supplement fig. 2C

JP-C protein sequence

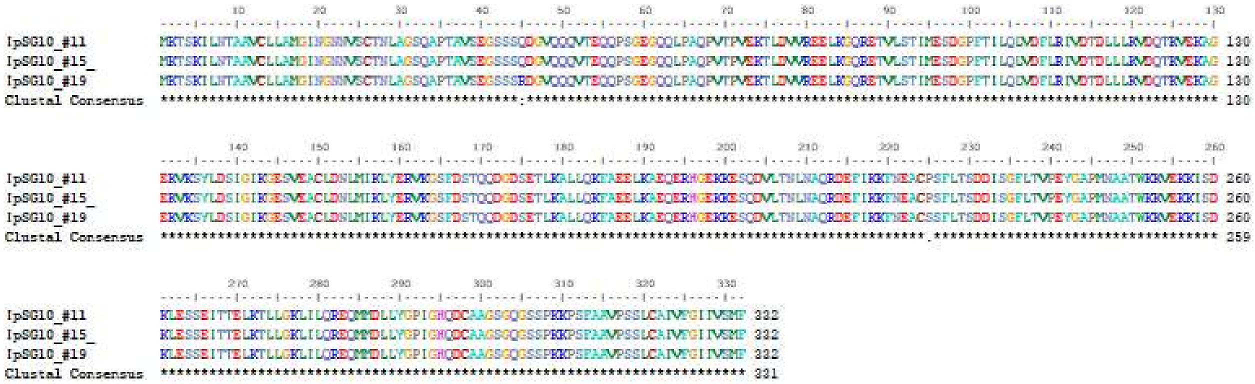

Supplement fig. 2D

EU-A1, EU-A2 protein sequence

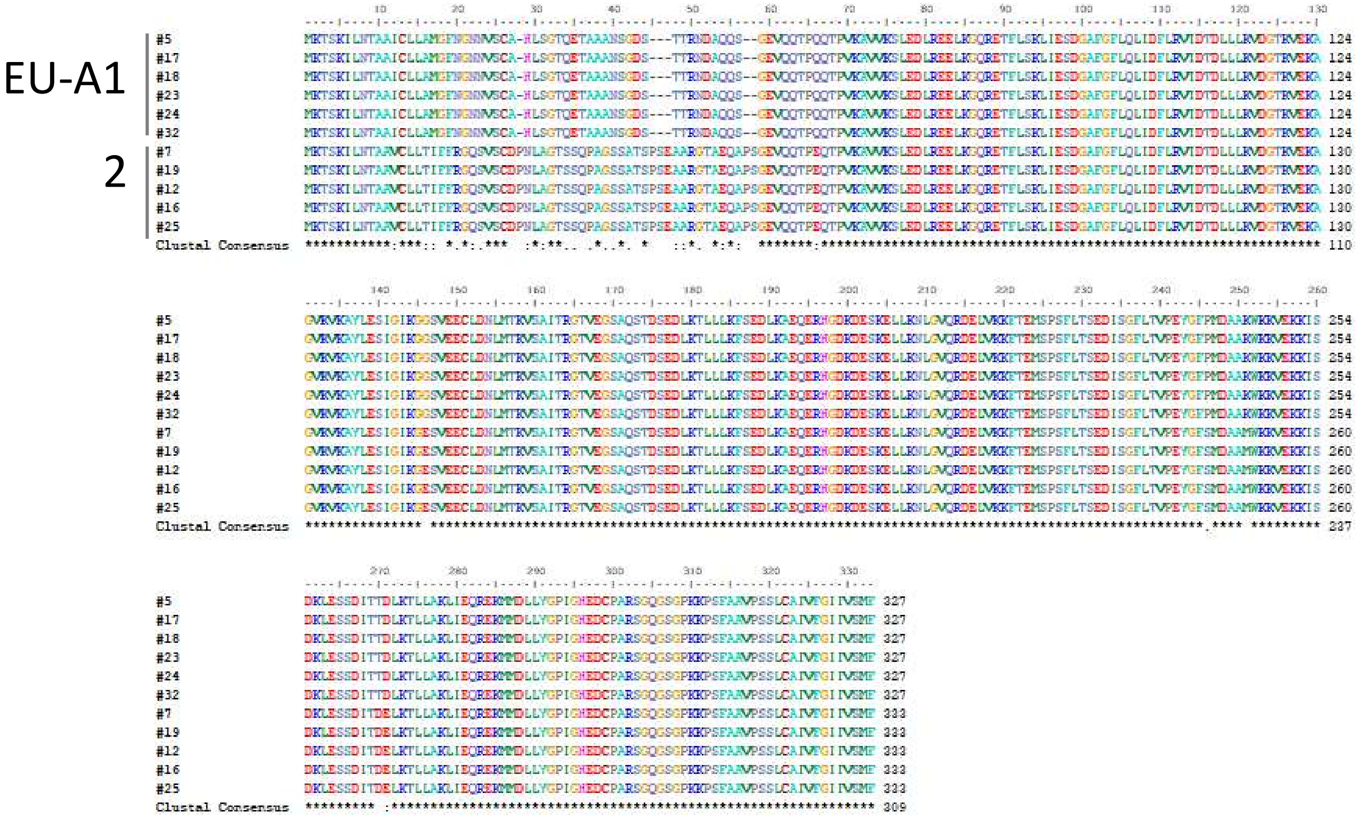

#### Supplement fig. 3A

### Deer JPA

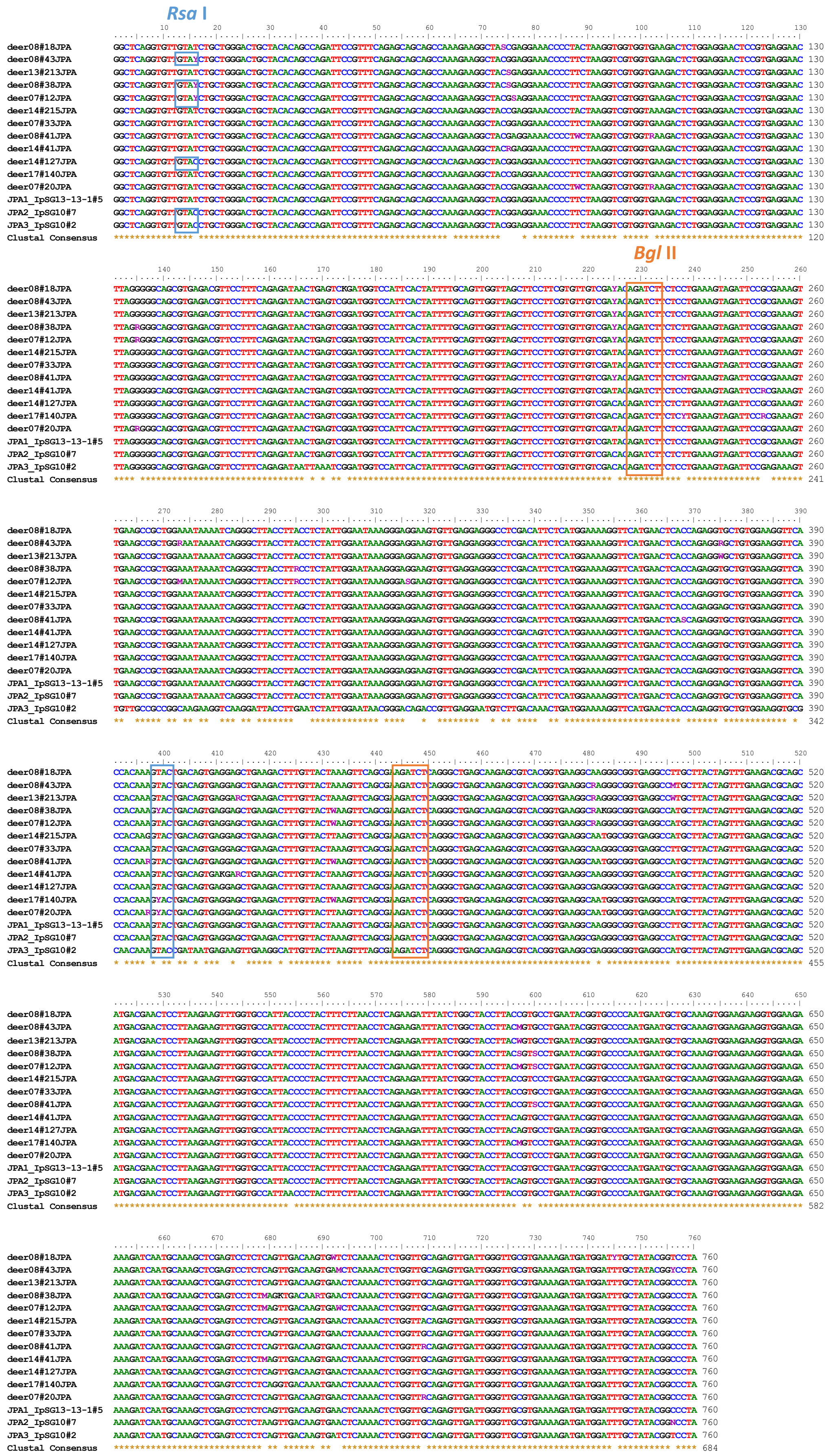

Supplement fig. 3A cont.

Deer JPA (amino acid sequence)

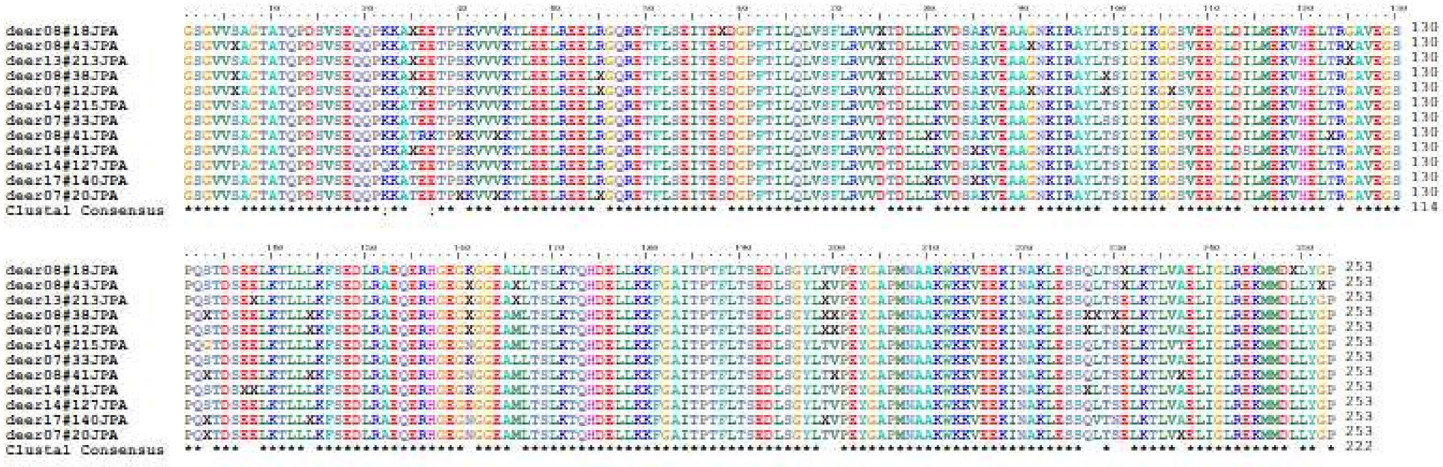

Supplement fig. 3B

Deer JPB

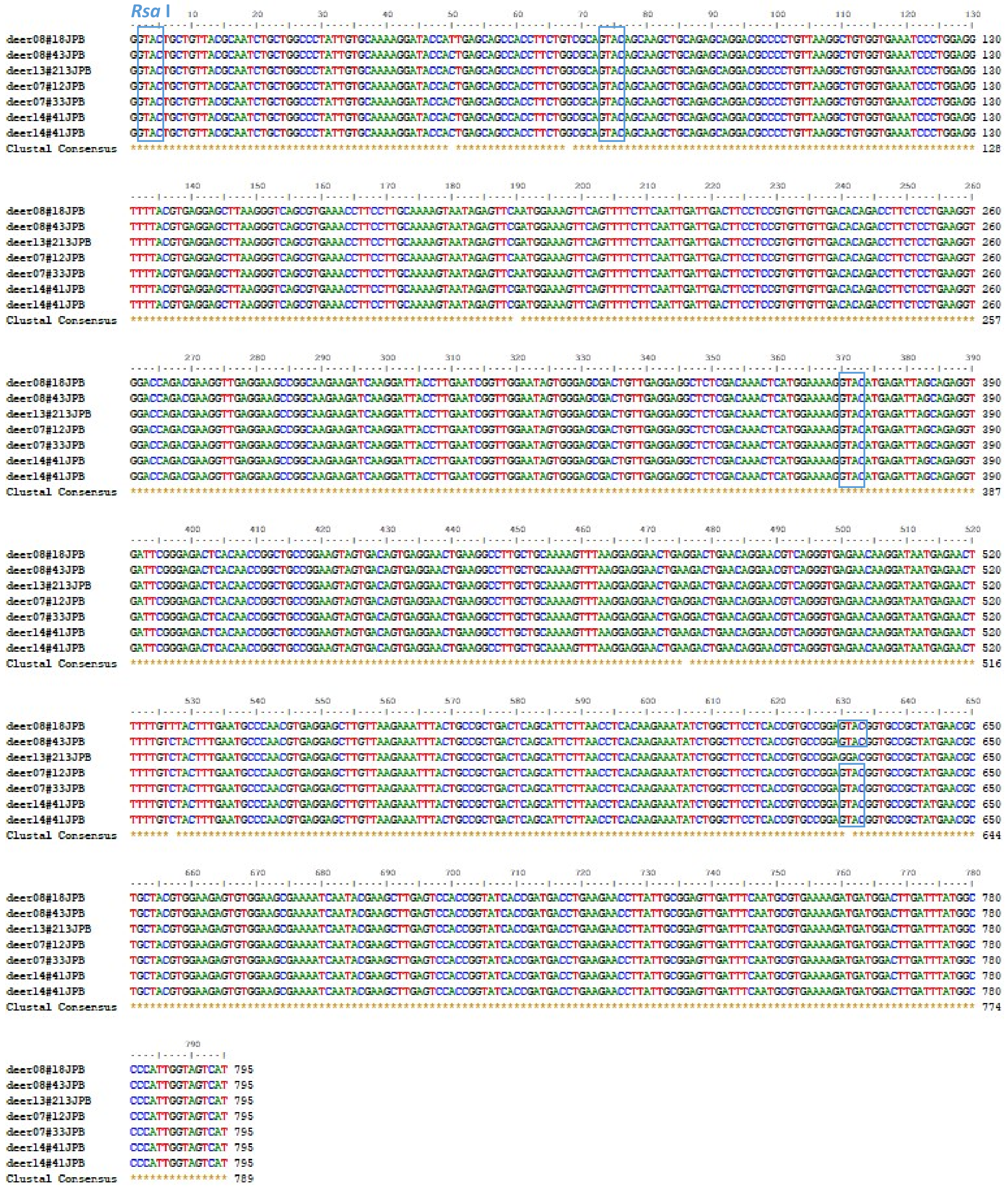

### Supplement fig. 3B cont.

#### Deer JPB (amino acid sequence)

```
                            10        20        30        40        50        60        70        80        90       100       110       120       130
                    ....|....|....|....|....|....|....|....|....|....|....|....|....|....|....|....|....|....|....|....|....|....|....|....|....|....|
deer08#18JPB        GTAVTQSAGPIVQKDTIEQPPSVAVQQAAEQDAPVKAVVKSLEVLREELKGQRETFLAKVIEFNGKFSFLQLIDFLRVVDTDLLLKVDQTKVEEAGKKIKDYLESVGIVGATVEEALDKLMEKVHEISRG 130
deer08#43JPB        GTAVTQSAGPIVQKDTTEQPPSGAVQQAAEQDAPVKAVVKSLEVLREELKGQRETFLAKVIEFDGKFSFLQLIDFLRVVDTDLLLKVDQTKVEEAGKKIKDYLESVGIVGATVEEALDKLMEKVHEISRG 130
deer13#213JPB       GTAVTQSAGPIVQKDTTEQPPSGAVQQAAEQDAPVKAVVKSLEVLREELKGQRETFLAKVIEFDGKFSFLQLIDFLRVVDTDLLLKVDQTKVEEAGKKIKDYLESVGIVGATVEEALDKLMEKVHEISRG 130
deer07#12JPB        GTAVTQSAGPIVQKDTTEQPPSGAVQQAAEQDAPVKAVVKSLEVLREELKGQRETFLAKVIEFNGKFSFLQLIDFLRVVDTDLLLKVDQTKVEEAGKKIKDYLESVGIVGATVEEALDKLMEKVHEISRG 130
deer07#33JPB        GTAVTQSAGPIVQKDTTEQPPSGAVQQAAEQDAPVKAVVKSLEVLREELKGQRETFLAKVIEFNGKFSFLQLIDFLRVVDTDLLLKVDQTKVEEAGKKIKDYLESVGIVGATVEEALDKLMEKVHEISRG 130
deer14#41JPB        GTAVTQSAGPIVQKDTTEQPPSGAVQQAAEQDAPVKAVVKSLEVLREELKGQRETFLAKVIEFDGKFSFLQLIDFLRVVDTDLLLKVDQTKVEEAGKKIKDYLESVGIVGATVEEALDKLMEKVHEISRG 130
deer14#41JPB        GTAVTQSAGPIVQKDTTEQPPSGAVQQAAEQDAPVKAVVKSLEVLREELKGQRETFLAKVIEFDGKFSFLQLIDFLRVVDTDLLLKVDQTKVEEAGKKIKDYLESVGIVGATVEEALDKLMEKVHEISRG 130
Clustal Consensus   **************** ***** *****************************************:***************************************************************** 128

                           140       150       160       170       180       190       200       210       220       230       240       250       260
                    ....|....|....|....|....|....|....|....|....|....|....|....|....|....|....|....|....|....|....|....|....|....|....|....|....|....|
deer08#18JPB        DSGDSQPAAGSSDSEELKALLQKFKEELRTEQERQGENKDNENFLFTLNAQREELVKKFTAADSAFLTSQEISGFLTVPEYGAAMNAATWKSVEAKINTKLESTGITDDLKNLIAELISMREKMMDLIYG 260
deer08#43JPB        DSGDSQPAAGSSDSEELKALLQKFKEELKTEQERQGENKDNENFLSTLNAQREELVKKFTAADSAFLTSQEISGFLTVPEYGAAMNAATWKSVEAKINTKLESTGITDDLKNLIAELISMREKMMDLIYG 260
deer13#213JPB       DSGDSQPAAGSSDSEELKALLQKFKEELKTEQERQGENKDNENFLSTLNAQREELVKKFTAADSAFLTSQEISGFLTVPEDGAAMNAATWKSVEAKINTKLESTGITDDLKNLIAELISMREKMMDLIYG 260
deer07#12JPB        DSGDSQPAAGSSDSEELKALLQKFKEELRTEQERQGENKDNENFLSTLNAQREELVKKFTAADSAFLTSQEISGFLTVPEYGAAMNAATWKSVEAKINTKLESTGITDDLKNLIAELISMREKMMDLIYG 260
deer07#33JPB        DSGDSQPAAGSSDSEELKALLQKFKEELRTEQERQGENKDNENFLSTLNAQREELVKKFTAADSAFLTSQEISGFLTVPEYGAAMNAATWKSVEAKINTKLESTGITDDLKNLIAELISMREKMMDLIYG 260
deer14#41JPB        DSGDSQPAAGSSDSEELKALLQKFKEELKTEQERQGENKDNENFLSTLNAQREELVKKFTAADSAFLTSQEISGFLTVPEYGAAMNAATWKSVEAKINTKLESTGITDDLKNLIAELISMREKMMDLIYG 260
deer14#41JPB        DSGDSQPAAGSSDSEELKALLQKFKEELKTEQERQGENKDNENFLSTLNAQREELVKKFTAADSAFLTSQEISGFLTVPEYGAAMNAATWKSVEAKINTKLESTGITDDLKNLIAELISMREKMMDLIYG 260
Clustal Consensus   ****************************:***************** ********************************** ************************************************ 256

                    ....|
deer08#18JPB        PIGSH 265
deer08#43JPB        PIGSH 265
deer13#213JPB       PIGSH 265
deer07#12JPB        PIGSH 265
deer07#33JPB        PIGSH 265
deer14#41JPB        PIGSH 265
deer14#41JPB        PIGSH 265
Clustal Consensus   ***** 261
```

Supplement fig. 3C

### Deer JPC

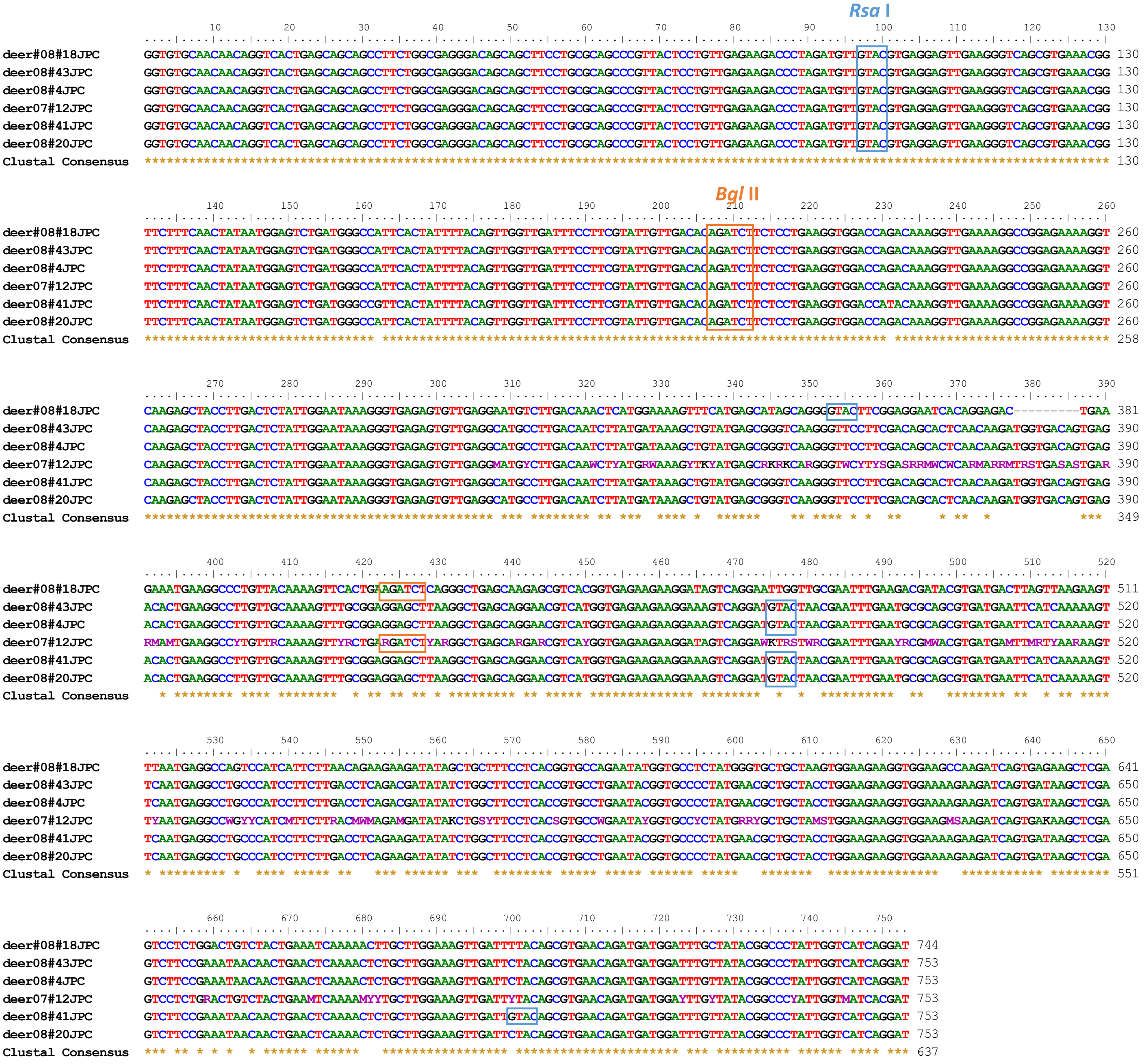

### Supplement fig. 3C

#### Deer JPC (amino acid sequence)

```
                           10        20        30        40        50        60        70        80        90       100       110       120       130
deer08#43          GVQQQVTEQQPSGEGQQLPAQPVTPVEKTLDVVREELKGQRETVLSTIMESDGPFTILQLVDFLRIVDTDLLLKVDQTKVEKAGEKVKSYLDSIGIKGESVEACLDNLMIKLYERVKGSFDSTQQDGDSE 130
deer08#4           GVQQQVTEQQPSGEGQQLPAQPVTPVEKTLDVVREELKGQRETVLSTIMESDGPFTILQLVDFLRIVDTDLLLKVDQTKVEKAGEKVKSYLDSIGIKGESVEACLDNLMIKLYERVKGSFDSTQQDGDSE 130
deer08#41          GVQQQVTEQQPSGEGQQLPAQPVTPVEKTLDVVREELKGQRETVLSTIMESDGPFTILQLVDFLRIVDTDLLLKVDHTKVEKAGEKVKSYLDSIGIKGESVEACLDNLMIKLYERVKGSFDSTQQDGDSE 130
deer08#20          GVQQQVTEQQPSGEGQQLPAQPVTPVEKTLDVVREELKGQRETVLSTIMESDGPFTILQLVDFLRIVDTDLLLKVDQTKVEKAGEKVKSYLDSIGIKGESVEACLDNLMIKLYERVKGSFDSTQQDGDSE 130
deer08#18          GVQQQVTEQQPSGEGQQLPAQPVTPVEKTLDVVREELKGQRETVLSTIMESDGPFTILQLVDFLRIVDTDLLLKVDQTKVEKAGEKVKSYLDSIGIKGESVEECLDKLMEKFHEHSRGTSEESQ---ETE 127
deer07#12          GVQQQVTEQQPSGEGQQLPAQPVTPVEKTLDVVREELKGQRETVLSTIMESDGPFTILQLVDFLRIVDTDLLLKVDQTKVEKAGEKVKSYLDSIGIKGESVEXXLDXXMXKXXEXXXGXXXXXXXXXXXX 130
Clustal Consensus  ****************************************************************************:*************************  **  * *  *   *             108

                          140       150       160       170       180       190       200       210       220       230       240       250
deer08#43          TLKALLQKFAEELKAEQERHGEKKESQDVLTNLNAQRDEFIKKFNEACPSFLTSDDISGFLTVPEYGAPMNAATWKKVEKKISDKLESSEITTELKTLLGKLILQREQMMDLLYGPIGHQD 251
deer08#4           TLKALLQKFAEELKAEQERHGEKKESQDVLTNLNAQRDEFIKKFNEACPSFLTSDDISGFLTVPEYGAPMNAATWKKVEKKISDKLESSEITTELKTLLGKLILQREQMMDLLYGPIGHQD 251
deer08#41          TLKALLQKFAEELKAEQERHGEKKESQDVLTNLNAQRDEFIKKFNEACPSFLTSEDISGFLTVPEYGAPMNAATWKKVEKKISDKLESSEITTELKTLLGKLIVQREQMMDLLYGPIGHQD 251
deer08#20          TLKALLQKFAEELKAEQERHGEKKESQDVLTNLNAQRDEFIKKFNEACPSFLTSEDISGFLTVPEYGAPMNAATWKKVEKKISDKLESSEITTELKTLLGKLILQREQMMDLLYGPIGHQD 251
deer08#18          EMKALLQKFTEDLRAEQERHGEKKDSQELVANLKTIRDDLVKKFNEASPSFLTEEDIAAFLTVPEYGASMGAAKWKKVEAKISEKLESSGLSTEIKNLLGKLILQREQMMDLLYGPIGHQD 248
deer07#12          XXKAXXQKXXXDXXAEXXRXGEKKDSQXXXXNLXXXRDXXXXKXNEAXXXFXXXXDIXXFLXVXEXGAXMXAAXWKKVEXKISXKLESSXLSTEXKXXLGKLIXQREQMMXLXYGXIGXHD 251
Clustal Consensus    **  **   :  **  * ****:**    **   **    * ***   *    **  ** * * ** * ** ***** *** ***** ::** *  ***** ****** * ** ** :*
```

Supplement fig. 3D

#### Deer07#12 JPC

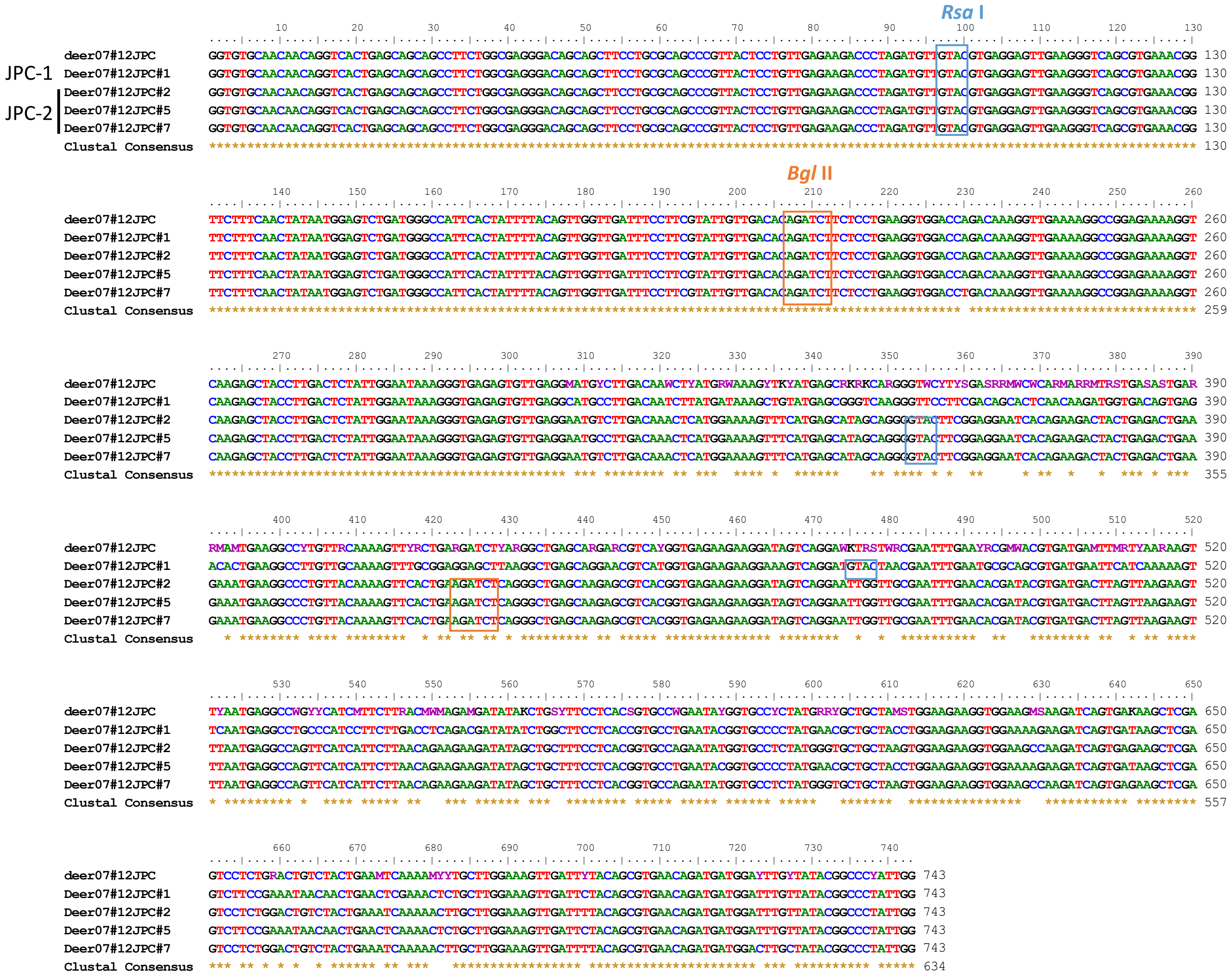


#### Deer07#12 JPC (protein sequence)

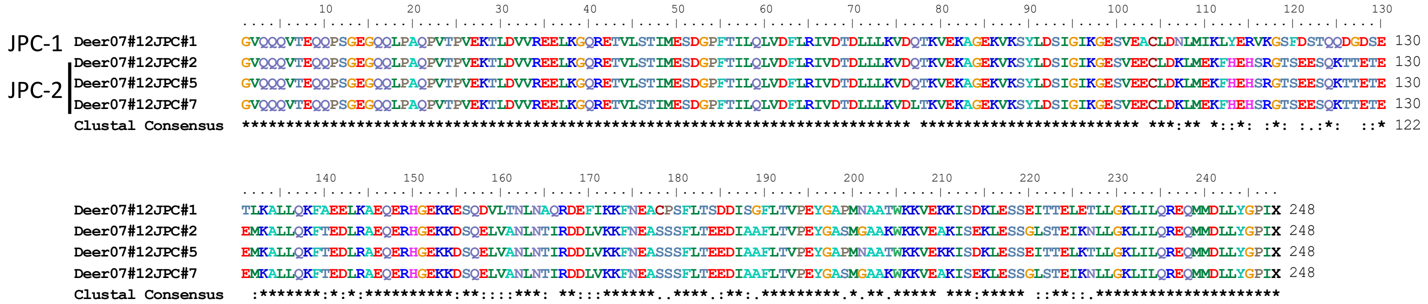
